# Oncoprotein CoAA repeats interact with RNA polymerase II CTD repeats

**DOI:** 10.1101/671156

**Authors:** Shiqin Xiong, Yang S. Brooks, Zheqiong Yang, Jiacai Wu, Liyong Zhang, William S. Dynan, Wei Xu, Bert W. O’Malley, Lan Ko

**Author notes:** Address correspondence to: Lan Ko, MD/PhD, Augusta, GA, USA.

## Abstract

The heptad repeating sequence of the C-terminal domain (CTD) of the largest subunit of RNA polymerase II is highly conserved in eukaryotes. In yeast, a CTD code consisting of pairs of heptad repeats is essential for viability. However, the strict requirement of diheptad repeats for the CTD function in transcription and splicing is unexplained. Here we show that CoAA (gene symbol RBM14), an oncoprotein and mammalian transcriptional coactivator, possesses diheptad repeats and directly interacts with the CTD. CoAA comprises 27 copies of tyrosine-rich repeats and regulates pre-mRNA synthesis and alternative splicing. Tyrosine substitutions in either the CoAA repeats or the CTD repeats diminish their interactions. Ser2- or Ser5-phosphorylated CTD peptides exhibit higher binding affinity to CoAA than the corresponding non-phosphorylated CTD peptide. CoAA dynamically interacts with both the CTD and hnRNP M, which is an alternative splicing regulator also comprising diheptad repeats. Arginine methylation of CoAA switches its interaction from the hnRNP M repeats to the CTD repeats. This study provides a mechanism for CoAA at the interface of transcription and alternative splicing, and explains the functional requirement of diheptad repeats in the CTD. In the human genome, tyrosine-rich repeats similar to the CoAA repeats were only found in six oncoproteins including EWS and SYT. We suggest that the diheptad sequence is one of the signature features for the CTD interaction among oncoproteins involved in transcription and alternative splicing. We anticipate that direct RNA Pol II interaction is a mechanism in oncogenesis.

## INTRODUCTION

The CTD of RNA polymerase II (RNAP II) largest subunit is required for pre-mRNA synthesis and RNA processing (1–4). This coordination requires the binding of an array of protein factors to the heptad sequence repeats (YSPTSPS) present in the CTD. Crystal structures have been determined in a number of CTD-modifying enzymes and factors involved in mRNA capping or polyadenylation (1). However, the puzzling requirement for multiple repeats in the CTD has yet to be explained. Extensive evidence supports for a requirement of tyrosine or serine phosphorylation and dephosphorylation of the CTD during transcription cycle (5–12). In addition, genetic studies in budding yeast demonstrate that pairs of heptad CTD repeats constitute the essential functional unit (13,14). Yeast survival relies on the presence of undisrupted diheptad repeats, in which a dual requirement of tyrosine and serine residues confers essential CTD activities. These observations predict that certain mammalian CTD-binding proteins will recognize its tyrosines in a diheptad pattern and its phosphorylated serines. These CTD-binding proteins shall be critical in transcription and alternative splicing.

### RESULTS AND DISCUSSION

Transcriptional coactivators stimulate gene activation and recruit the RNAP II complex during transcription (15). We find here that a transcriptional coactivator CoAA (<u>coa</u>ctivator activator, gene symbol RBM14) is a direct CTD-interacting protein with tandem diheptad repeats. CoAA was initially identified as a coactivator (16), and regulates transcription-coupled pre-mRNA alternative splicing (17). The human CoAA gene is amplified in cancers with recurrent deletion of its enhancer sequence (18). This deletion causes a defect in CoAA alternative splicing and blocks stem cell differentiation in tumorigenesis (19,20). CoAA gene amplification is found in mutant immune cells in tumor microenvironment of solid cancers associated with chronic inflammation (21).

CoAA contains two N-terminal RNA recognition motifs (RRMs) and a C-terminal activation domain containing 27 copies of tyrosine- and glutamine-rich repetitive sequence (Fig. 1a), previously termed the YxxQ repeats (16,18). The transcriptional activity of CoAA requires its YxxQ repeats. In the mammalian genome, repetitive sequences homologous to the YxxQ repeats of CoAA are present in only six oncoproteins including SYT and the EWS family members based on the database analysis at ScanProsite (18,22) (Supplementary Fig. S1). The sequence repeats in these oncoproteins are transcriptionally active and have been shown to be necessary for tumor initiation (21,23).

**Figure 1.**
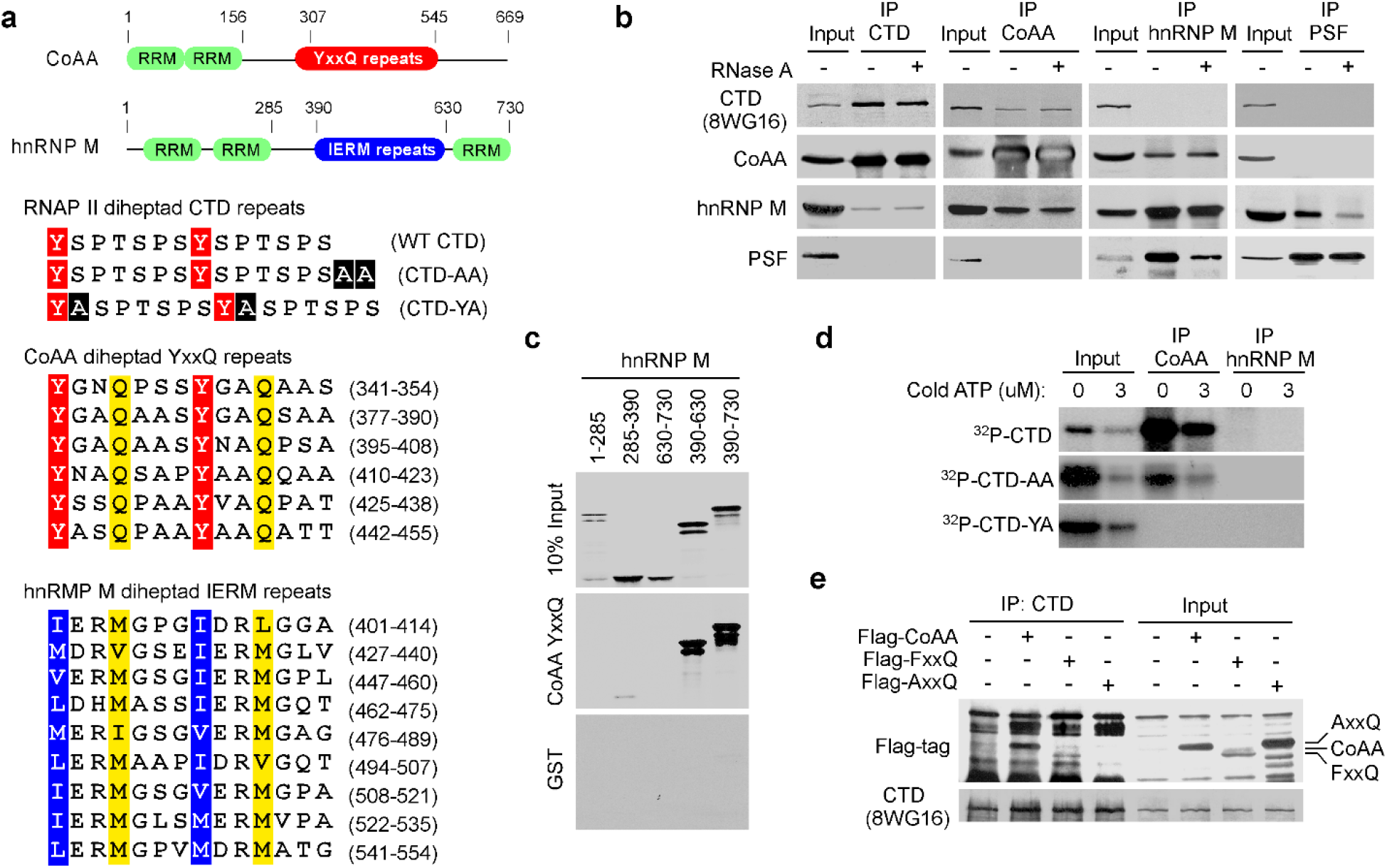
CoAA interacts with the repeating sequences of hnRNP M and the CTD of RNAP II. **a**, Protein domains of CoAA and hnRNP M with numbers indicating amino acids. The pairs of heptad repeats are highlighted with conserved tyrosine, glutamine, and hydrophobic residues in the CTD, CoAA, and hnRNP M (24). Alanine insertions in CTD-AA and CTD-YA are highlighted in black. **b**, Coimmunoprecipitation of HeLa nuclear extracts in the presence or absence of RNase A using antibodies as indicated. **c**, *In vitro* binding assay using GST-YxxQ (307-545) and *in vitro*-translated ^35^S-methionine-labeled hnRNP M fragments. **d**, GST-CTD and its mutants were ^32^P-γ-ATP-labeled by Cdc2 kinase and incubated with immunoprecipitated CoAA or hnRNP M. Bound proteins were detected by autoradiography. **e**, Coimmunoprecipitation of Flag-tagged CoAA or its mutants AxxQ and FxxQ and RNAP II in 293 cells.

To seek proteins that interact with the intriguing YxxQ repeats of CoAA, we carried out mass spectrometry using recombinant YxxQ repeats (307-545) as bait. The most efficiently bound proteins were identified as heterogeneous ribonucleoprotein hnRNP M isoforms (Supplementary Table S1, in a separate PDF file), and immunoblotting of bound proteins also detected RNAP II (Supplementary Fig. S2). hnRNP M is known to interact with pre-mRNA and to regulate pre-mRNA splicing on/off during heat shock (24–26). hnRNP M contains two N-terminal RRM domains, 27 copies of methionine-rich IERM repeats (24), and a C-terminal RRM domain (Fig. 1a and Supplementary Fig. S3). CoAA has a related secondary structure with hnRNP M and also belongs to the hnRNP family (16). Due to less abundance, CoAA protein was not initially identified in the hnRNP protein complex relying 2D gel analysis. The presence of the imperfect 27-copy repeats in both CoAA and hnRNP M is a striking shared feature in addition to their RRM domains. Importantly, pairs of heptads were identified in both the YxxQ and the IERM repeats similar to that of the CTD (Fig. 1a and Supplementary Fig. S4).

Both CoAA and hnRNP M are RRM proteins involved in transcription and alternative splicing. We next investigated their RNA-dependent *in vivo* associations with transcription and splicing complexes using RNAP II and the splicing regulator PSF as markers, respectively. Using coimmunoprecipitation (co-IP) of nucleus lysates from cells, CoAA interacted with hnRNP M; however, their association towards transcription and splicing complexes appeared to be dynamic. Specifically, CoAA is closely associated with RNAP II but not PSF, independent of RNA. hnRNP M is closely associated with PSF but not RNAP II, dependent of RNA (Fig. 1b). Consistently, CoAA but not hnRNP M is found in the complex containing phosphorylated RNAP II (Supplementary Fig. S5), and only a fraction of CoAA is colocalized with RNAP II in the nucleus (Supplementary Fig. S6). The interaction of CoAA and hnRNP M may reflect a dynamic association of transcription and splicing complexes. These results nonetheless showed that different pools of CoAA and hnRNP M exist *in vivo*.

To test the potential direct interactions among repeat-containing CoAA, hnRNP M, and RNAP II, we carried out *in vitro* pull-down studies with mutagenesis. CoAA YxxQ repeats directly interacted with hnRNP M IERM repeats *in vitro* (Fig. 1c). In addition, consistent with co-IP observations, CoAA but not hnRNP M interacted with serine-phosphorylated recombinant CTD (Fig. 1d). When the CTD was mutated with alanine insertions (CTD-YA) to disrupt its diheptad feature (Fig. 1a), a lethal phenotype in yeast (14), the CTD binding to CoAA was diminished (Fig. 1d). When the CTD was mutated but preserving the diheptad repeats (CTD-AA), a viable phenotype in yeast, the CTD binding to CoAA was present except reduced. When mutations were introduced to the 27 tyrosine residues in CoAA YxxQ repeats through gene synthesis, alanine mutations AxxQ abolished the CTD binding but phenylalanine mutations FxxQ did not (Fig. 1e). Together, these data indicate that the tyrosine residues in CoAA are required for the diheptad CTD interaction.

We further confirmed the direct interaction between CoAA and the CTD through peptide binding studies. Recombinant GST fusion CoAA YxxQ repeats (307-545) were coated on a 96-well plate and incubated with synthetic biotin-labeled CTD diheptad peptides. Bound peptides were detected via streptavidin conjugates. EC_50_ can be measured using binding kinetics. We compared unphosphorylated, Ser2- or Ser5-phosphorylated and tyrosine to alanine mutated CTD peptides. The Ser2- and Ser5-phosphorylated diheptad peptides bound to the YxxQ repeats with median effective concentration EC_50_ at 135 nM and 101 nM, respectively (Fig. 2a). The binding of unphosphorylated CTD peptide occurred at a much lower affinity with EC_50_ at 1066 nM. However, the tyrosine to alanine mutant of the CTD peptides essentially abolished the interaction of the YxxQ repeats, binding was not detectable (ND). As controls, the IERM peptide bound to the YxxQ repeats with EC_50_ at 83 nM. The recombinant CoAA AxxQ mutant failed to bind to any of these peptides (Fig. 2a and Supplementary Fig. S7). A reciprocal binding assay using the YxxQ diheptad peptide and recombinant mutated CTD-AA and CTD-YA proteins indicated the diheptad feature of the CTD is required for the interaction (Fig. 2b).

**Figure 2.**
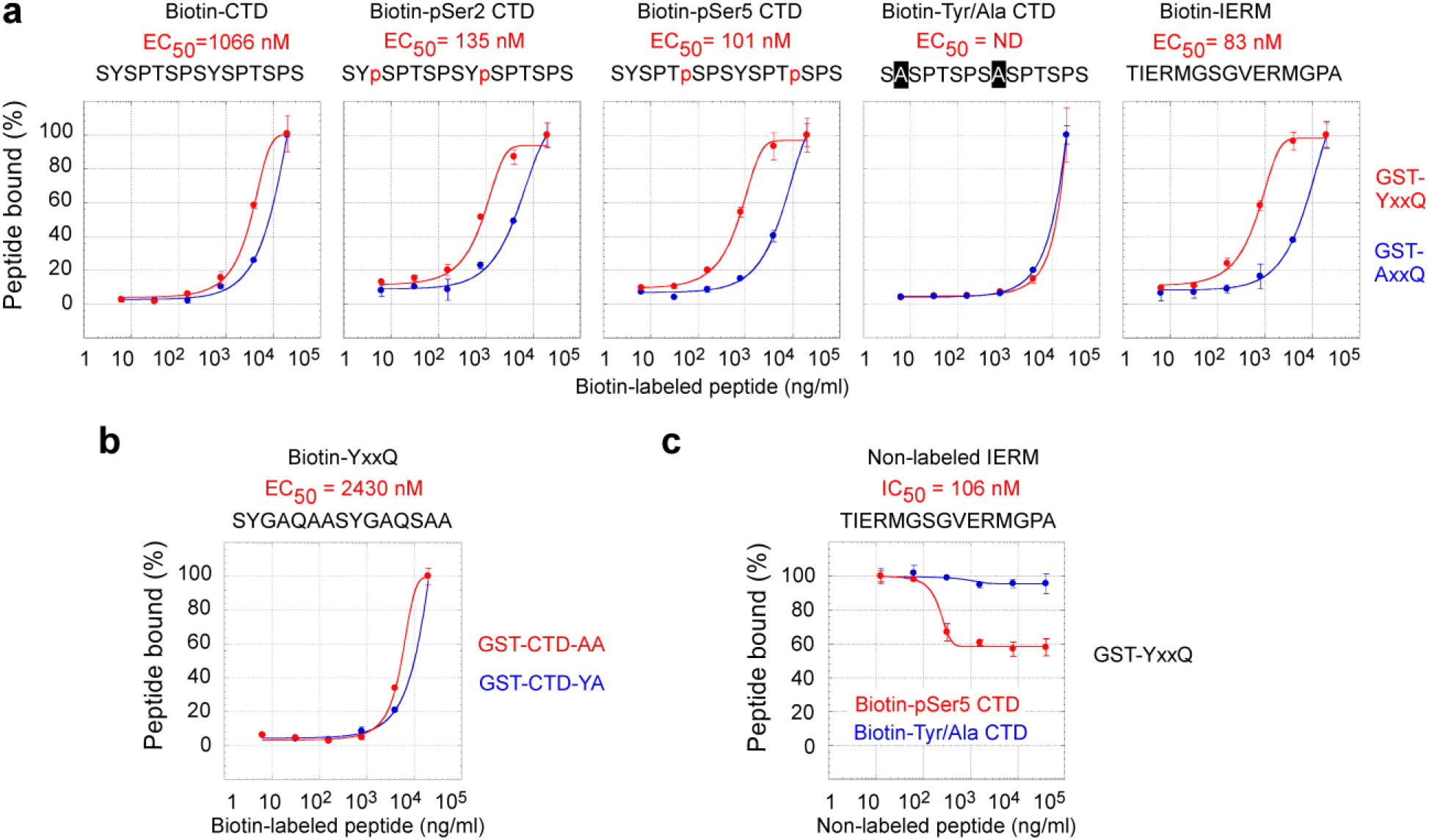
The CoAA YxxQ repeats recognize the tyrosine residues of serine phosphorylated diheptad CTD repeats. **a**, Recombinant GST-YxxQ and GST-AxxQ (307-545) were coated on 96-well plate. Biotin-labeled synthetic diheptad peptides, with sequences depicted, were incubated with increasing concentrations at 6.4, 32, 160, 800, 4000, and 20000 ng/ml. Bound biotin peptides were detected by streptavidin-HRP conjugates. EC_50_ values were determined by sigmoidal nonlinear regression curve fit. Only detectable EC_50_ values for YxxQ are shown. **b**, The binding of mutants GST-CTD-AA and GST-CTD-YA to the YxxQ diheptad peptide. **c**, The competition of non-labeled IERM peptide (12.8, 64, 320, 1600, 8000, 40000 ng/ml) with the binding of biotin-labeled Ser5-phosphorylated or Tyr/Ala mutated CTD peptides (4000 ng/ml) to GST-YxxQ. Data shown are means of duplicates ± s.e.m.

Furthermore, a competition assay showed that the IERM peptide was able to compete with the Ser5-phosphorylated but not with alanine mutated CTD for binding to CoAA (Fig. 2c). These data collectively indicate that the serine phosphorylation of CTD is critical for high binding affinity. The tyrosine residues in the CTD, similar to that in CoAA, are essential for the interaction. The diheptad peptides from both repeats are sufficient as minimal binding units consistent with the previous finding in yeast (13). *In vivo*, however, higher binding efficiency would be expected with longer repeats. It might also be physiologically meaningful that CoAA binds to the CTD and hnRNP M with comparable affinity so that their binding could be interchangeable upon regulation.

The transcriptional and splicing activities of CoAA and hnRNP M were further analyzed to gain more insights into their interrelationship. CoAA is transcriptionally active and hnRNP M is not (Fig. 3a). The tyrosine to alanine AxxQ but not to phenylalanine FxxQ mutations in the YxxQ repeats abolished CoAA activity, which can be explained by the requirement of tyrosine residues in the CTD interaction (Fig. 1e). When individual domains were tested by Gal4-fusion system, only the YxxQ or FxxQ repeats had potent transcriptional activity reflecting their direct interactions with the CTD (Fig. 3b).

**Figure 3.**
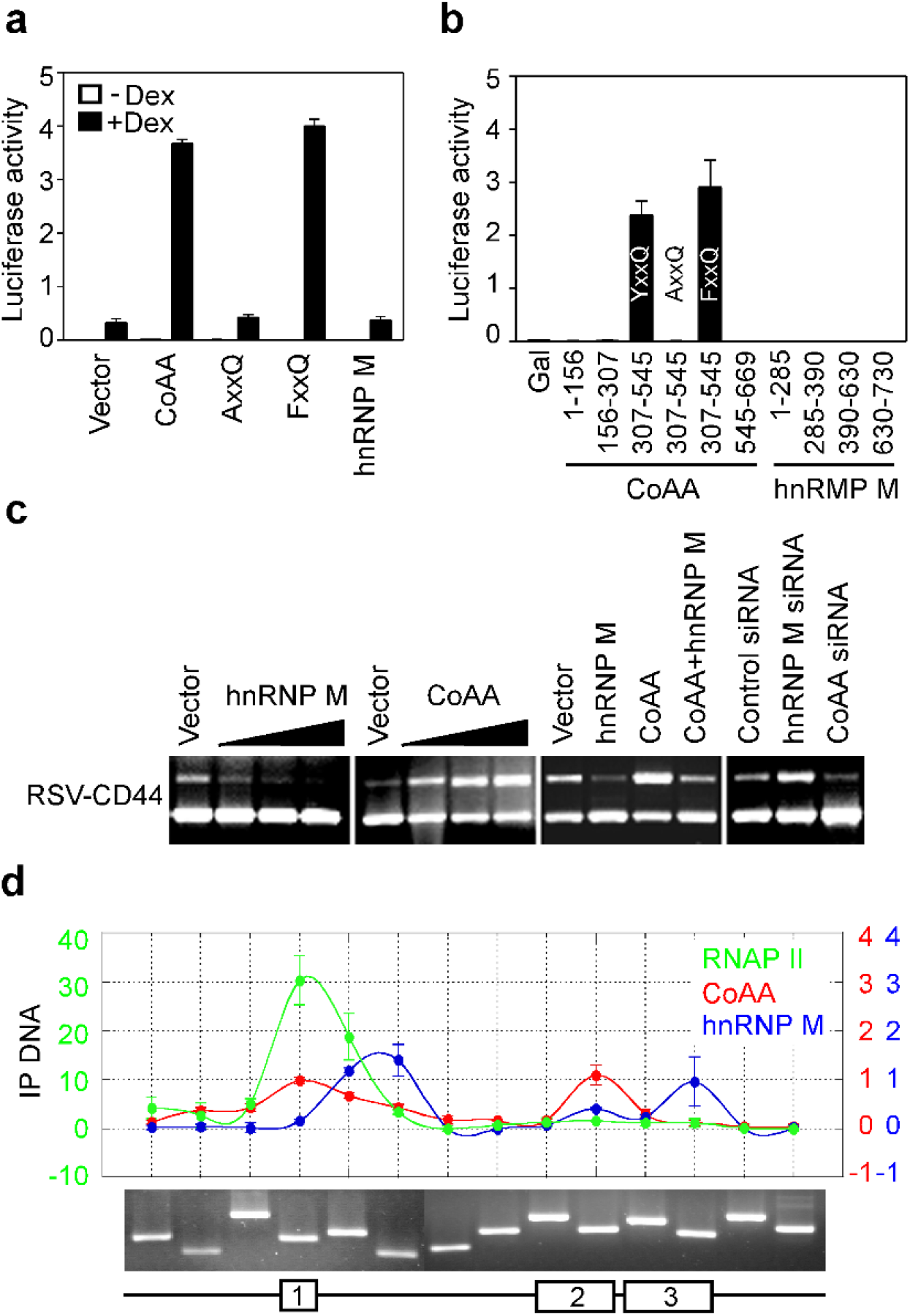
Transcriptional and splicing activities of CoAA and hnRNP M. **a**, Transcriptional activities of full length CoAA, mutants AxxQ or FxxQ, or hnRNP M were determined by transfection of CV-1 cells using a MMTV-luciferase reporter together with glucocorticoid receptor and induced by dexamethasone as ligand. **b**, Each protein domain of CoAA and hnRNP M as indicated were fused to Gal4 DNA-binding domain and tested for activity using a 5XGal-luciferase reporter. Luciferase activities shown are means of triplicate transfections ± s.e.m. **c**, A pETv5 splicing reporter containing alternatively spliced CD44 variable exon 5 was assayed by RT-PCR in 293 cells in the presence of overexpressed CoAA or hnRNP M (10, 50, 200 ng/well), or their combination (100 ng each), or their siRNA (100 nM). **d**, Chromatinimmunoprecipitation (ChIP) analysis of HeLa cells using antibodies against the CTD (8WG16), CoAA and hnRNP M. The human CoAA gene was used as template with its exons shown in the schematic diagram. Primer evaluation using human genomic DNA is shown below. Colored scales of y axis correspond to the graphs. Data shown are means of duplicates ± s.e.m. Primer sequences are listed in Supplementary Table S2.

Using a splicing cassette of CD44 minigene, we found that CoAA and hnRNP M counterregulate alternative splicing choices (Fig. 3c). In addition, chromatin immunoprecipitation (ChIP) analysis of the CoAA gene, known to be regulated by CoAA itself (18), showed non-overlapping chromatin binding profiles between CoAA and hnRNP M (Fig. 3d). While the chromatin occupancy of CoAA parallels with that of RNAP II at the CoAA exon regions, hnRNP M showed increased chromatin binding following the decrease of CoAA. CoAA and hnRNP M potentially balance exon inclusion and skipping through their heterodimerization. Their preferential association to RNAP II complex or splicing complex might provide a bridging connection for transcription-coupled splicing (Supplementary Fig. S4d), in which the higher transcription rate often links to one splicing choice and the lower rate to another (27).

Although CoAA and hnRNP M are likely regulated at multiple levels during transcription, we examined their protein arginine methylations for two reasons. First, the hnRNP family proteins are extensively methylated during transcription (28–30). Second, arginine dimethylation has been shown in EWS, an RRM protein with homology to the CoAA YxxQ repeats (31) (Supplementary Fig. S1). Our results indicated that CoAA but not hnRNP M is arginine methylated by CARM1 predominantly at the regions surrounding the YxxQ repeats (Fig. 4a and Supplementary Fig. S8a). When fractionated CoAA from cell nucleus was tested for the CTD interaction upon CARM1 methylation, CoAA that co-purified with hnRNP M had increased CTD binding preferentially to phosphorylated CTD. In contrast, CoAA free from hnRNP M failed to bind to the CTD (Fig. 4b-c). During dose-dependent CARM1 treatment, CoAA switched binding from hnRNP M to the CTD upon arginine methylation (Fig. 4d). Our results do not exclude the presence of additional possible regulations including methylation by other arginine methyltransferases, but nonetheless demonstrate regulated CoAA binding with hnRNP M switching to the CTD under methylation (Fig. 4e).

**Figure 4.**
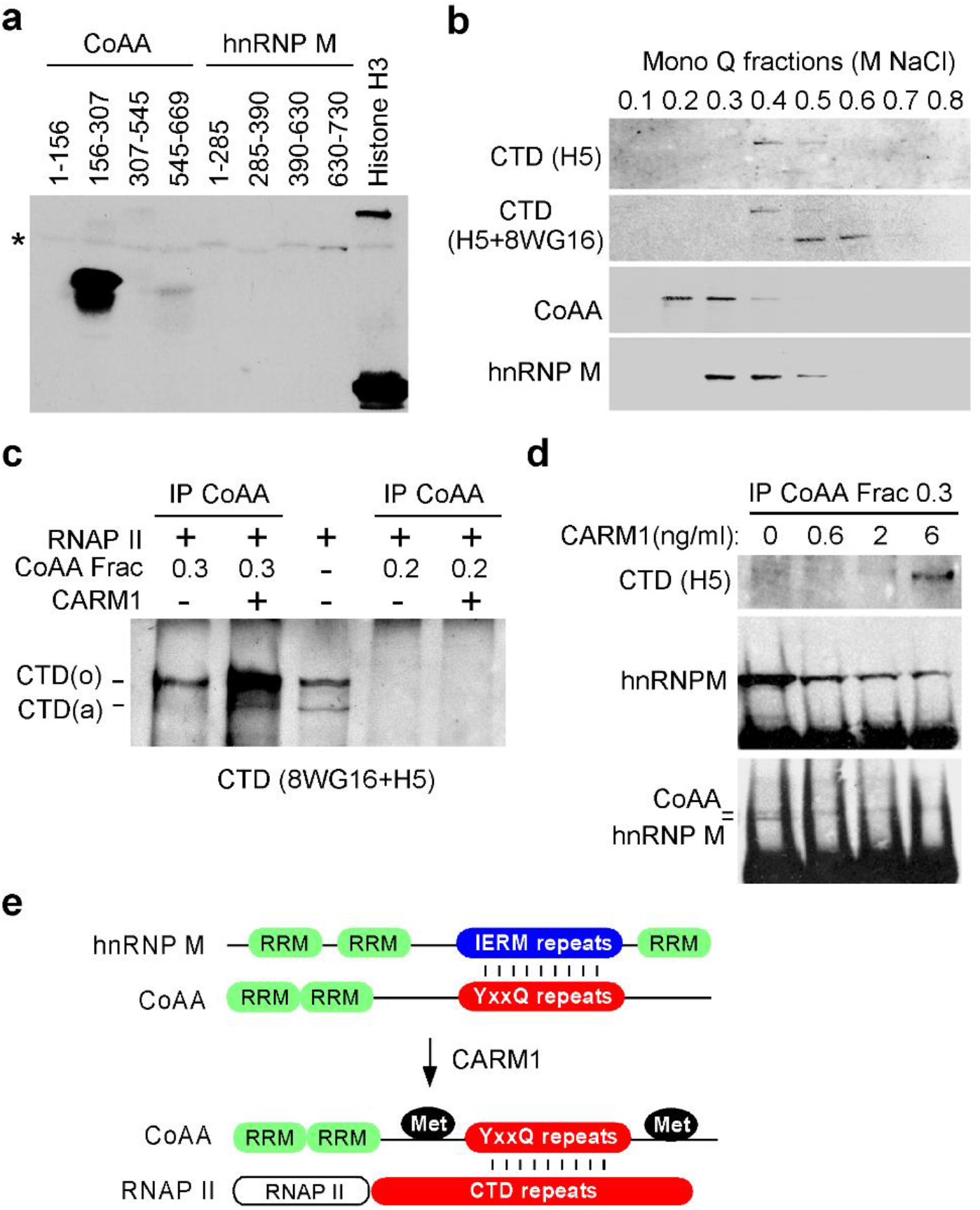
CoAA switches the interaction from hnRNP M to the CTD upon arginine methylation. **a**, GST fragments of CoAA or hnRNP M were arginine methylated by CARM1 methyltransferase and labeled with [^3^H]-AdoMet *in vitro* using Histone H3 as a positive control. Auto-methylated CARM1 is indicated by an asterisk (see Supplementary Fig. S8a for Coomassie blue staining of the gel). **b**, Partial purification of CoAA fractions by Mono Q ion exchange chromatography with step elution from 0.1 to 0.8 M NaCl. Each fraction was analyzed by Western blots using anti-CTD H5 (1:200), reprobed with 8WG16 (1:200), anti-CoAA, and anti-hnRNP M. **c**, Immunoprecipitated CoAA Mono Q fractions (0.2, 0.3 NaCl) were arginine methylated by CARM1 (6 ng/μl) *in vitro* before incubation with partially purified RNAP II derived from 0.4-0.5 M NaCl Mono Q fractions. Bound RNAP II was detected by combined anti-CTD antibodies 8WG16 and H5. **d**, CoAA from 0.3 M NaCl fraction was immunoprecipitated and methylated with increasing amounts of CARM1 (0. 0.6, 2, 6 ng/μl). The precipitates were incubated with the RNAP II fraction and the bound proteins were detected by Western blots using anti-CTD (H5), anti-hnRNP M, and anti-CoAA (re-probed using hnRNP M blot). **e**, Model of CoAA action. CoAA interacts with hnRNP M through their repeating sequences. Protein arginine methylation of CoAA by CARM1 induces the disassociation of hnRNP M and the association of the CTD of RNAP II.

The CTD heptad sequence is strongly conserved in eukaryotes. This study provides the first example to our knowledge of a CTD-binding protein with diheptad repeats. The CTD is a compact β-spiral structure and becomes relaxed or exposed upon serine phosphorylation (32). Consistent with this, CoAA recognizes the minimal CTD diheptad peptide through a pair of its tyrosine residues whose conformation is optimized by nearby Ser5 or Ser2 phosphorylation. On the other hand, the CoAA YxxQ repeats appear to require regulation, such as surrounding arginine methylations, in order to be accessible to the CTD. CoAA becomes more proteolytically sensitive upon arginine methylation as well as hnRNP M interaction (Supplementary Fig. S8b). Since all three molecules possess diheptad peptides, it remains to be determined whether hnRNP M interaction is required for CoAA prior to bind to the CTD. In conclusion, this study together indicates that CoAA diheptad repeats directly interacts with RNAP II CTD repeats.

The multiple tyrosine and glutamine-rich repeats in mammalian protein databases are restricted to only a few oncoproteins including CoAA (18). This implicates a fundamental important role of oncoproteins as the CTD interaction, whose defect impacts transcription-coupled alternative splicing. CoAA is previously shown to control stem cell differentiation at initial stage, and gene amplification in CoAA disrupts its own alternative splicing and blocks stem cell differentiation (18,19,33). Therefore, the defect in both stem cell regulation and RNA polymerase II interaction in oncoproteins is a conceivable mechanism in oncogenesis (21).

## ACKNOWLEDGEMENTS

We thank Dr. Dorothy Tuan for critical suggestions, and Dr. Yun Kyoung Kang for help. We thank Dr. John W. Stiller and Dr. Pengda Liu for providing CTD-AA and CTD-YA, Dr. Harald König for providing pET-v5 minigene, Dr. Michael Stallcup for providing CARM1, and Dr. James L. Manley for providing the GST-CTD construct. This work was supported in part by the Georgia Cancer Coalition (L.K.), the NIH (W.X.) and the NIH NIDDK (B.W.O.).

---

**Supplementary Figure S1.**
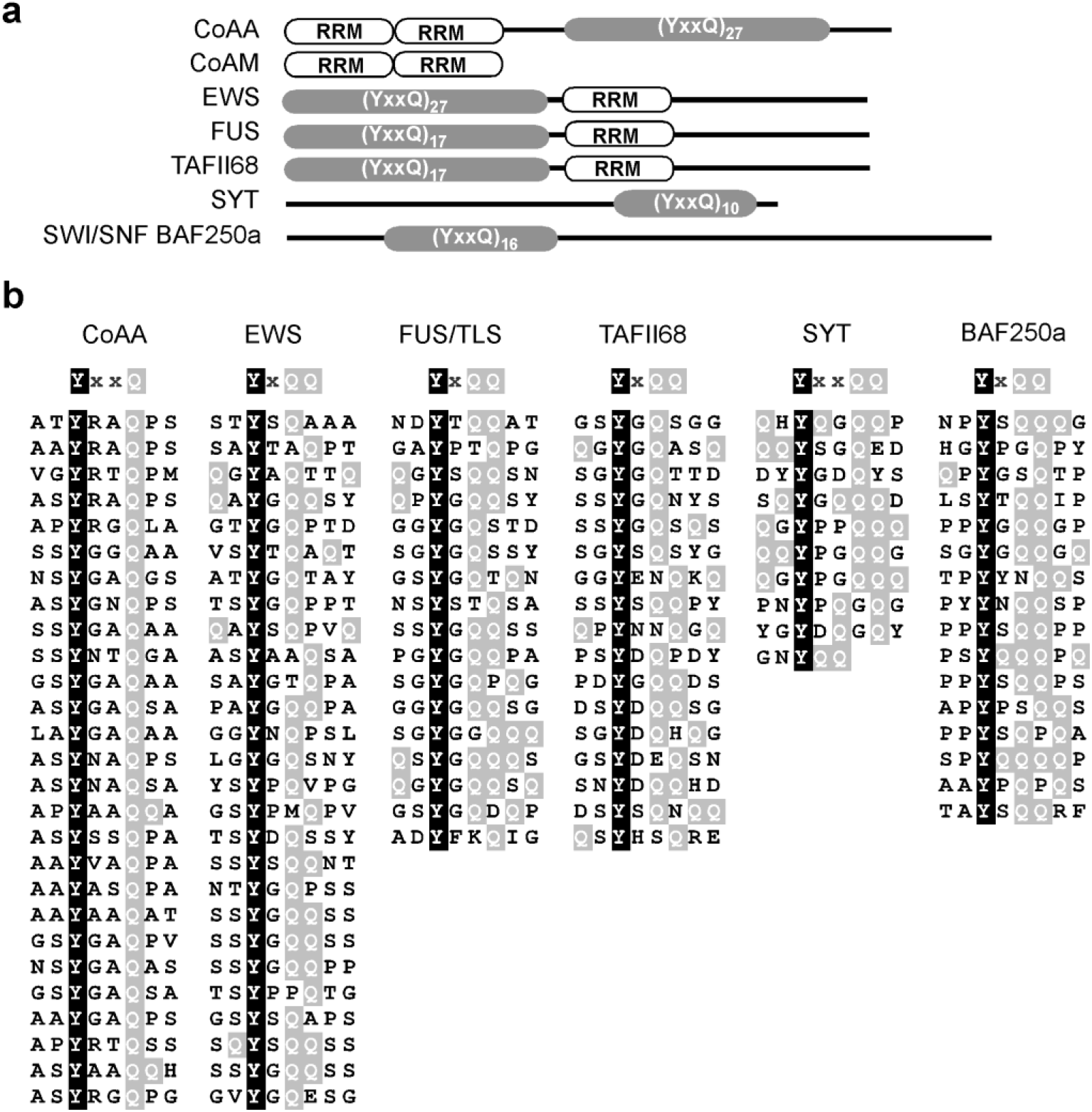
Oncoproteins with tyrosine-rich repeats. Swiss-Port and TrEMBL protein databases were analyzed with ScanProsite (http://us.expasy.org/tools/scanprosite/). Prosite format scanned: Y-X(1,2)-Q(1,2)-X(1,4)-Y-X(1,2)-Q(1,2)-X(1,4)-Y-X(1,2)-Q(1,2). The taxonomic species filters were set as *Homo sapiens, Mus musculus, Rattus norvegicus and Bos Taurus*. Sequences identified with more than three hits, *i.e.* nine copies of the YxxQ motifs, were selected and listed with their identities. **a**, Schematic diagrams of identified oncoproteins. RRM domains are indicated in white and YxxQ domains in grey. CoAA dominant negative splice variant CoAM is also shown. **b**, Alignment of YxxQ repeats. Tyrosine residues (Y) are shaded in black. Glutamine residues (Q) are shaded in grey. Additional two residues flanking the YxxQ motif are shown. In the human genome, CoAA is a unique molecule.

**Supplementary Figure S2.**
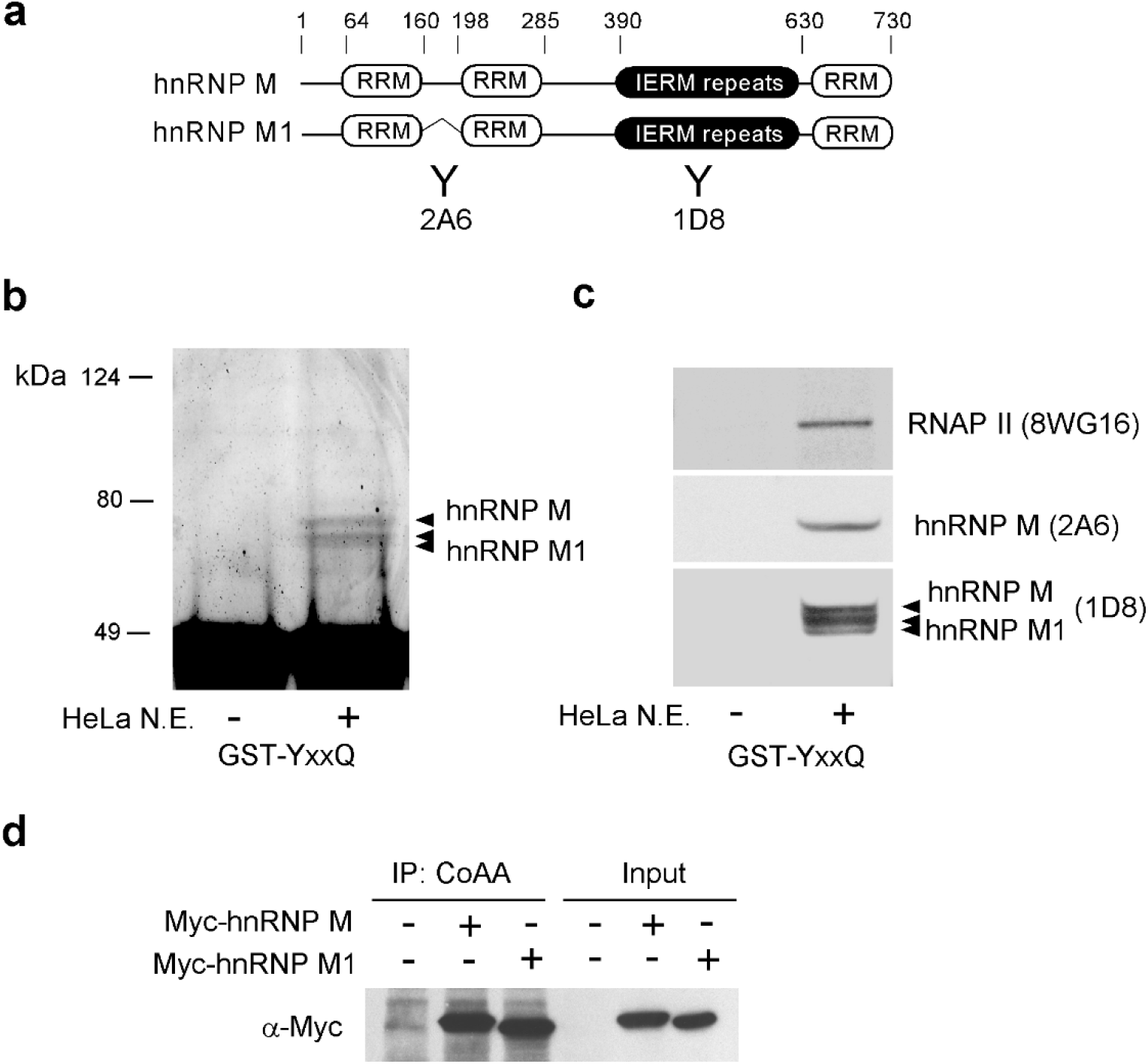
Identification of hnRNP M isoforms as CoAA-interacting proteins. **a**, Schematic representation of domain structures of hnRNP M and its alternatively spliced isoform hnRNP M1. Antibodies 2A6 and 1D8 are indicated. **b**, Recombinant CoAA protein GST-YxxQ (307-545) was incubated with HeLa nuclear extracts (N.E.). Bound proteins were resolved by SDS-PAGE followed by Coomassie blue staining. Proteins excised from the gel were identified by mass spectrometry as hnRNP M isoforms. **c**, Western blot analyses of bound hnRNP M isoforms and RNAP II. Multiple bands of hnRNP M and hnRNP M1 are known to be the results of phosphorylation status. **d**, *In vivo* interaction of CoAA and Myc-tagged hnRNP M isoforms was analyzed by co-immunoprecipitation in 293 cells.

**Supplementary Figure S3.**
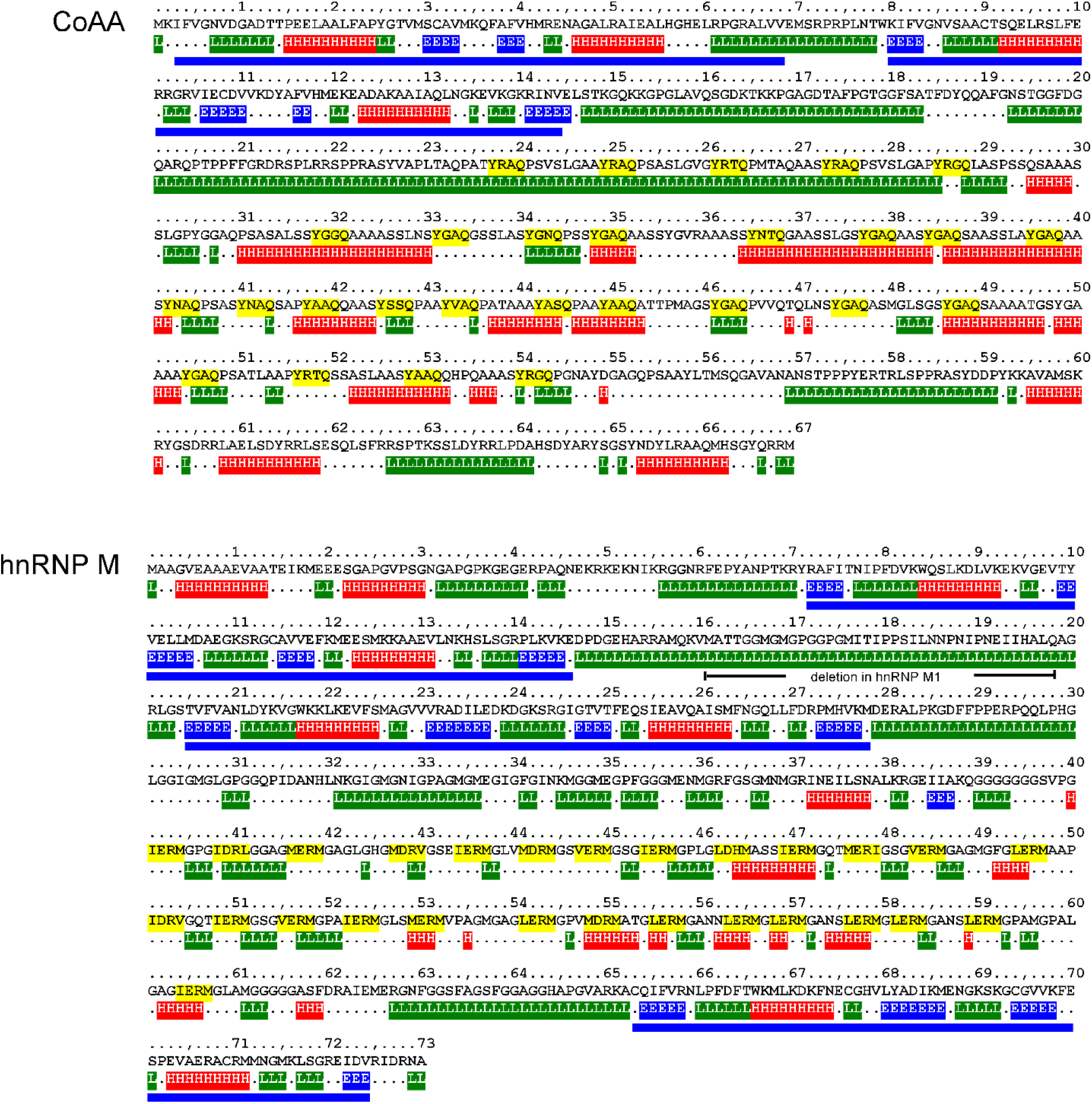
Predicted secondary structure of CoAA and hnRNP M. The protein secondary structure of CoAA and hnRNP M were analyzed by PHD (Profile Network Prediction HeiDelberg). Predicted α-helix (H, red), β-strand (E, blue) and unstructured loop region (L, green) are shown below the amino acid sequences, which are numbered at every 10th residue. The conserved RRM domains are indicated with blue bars. The YxxQ and IERM repeats are highlighted in yellow. The position of a natural deletion due to alternative splicing in hnRNP M1 is indicated with a bracket.

**Supplementary Figure S4.**
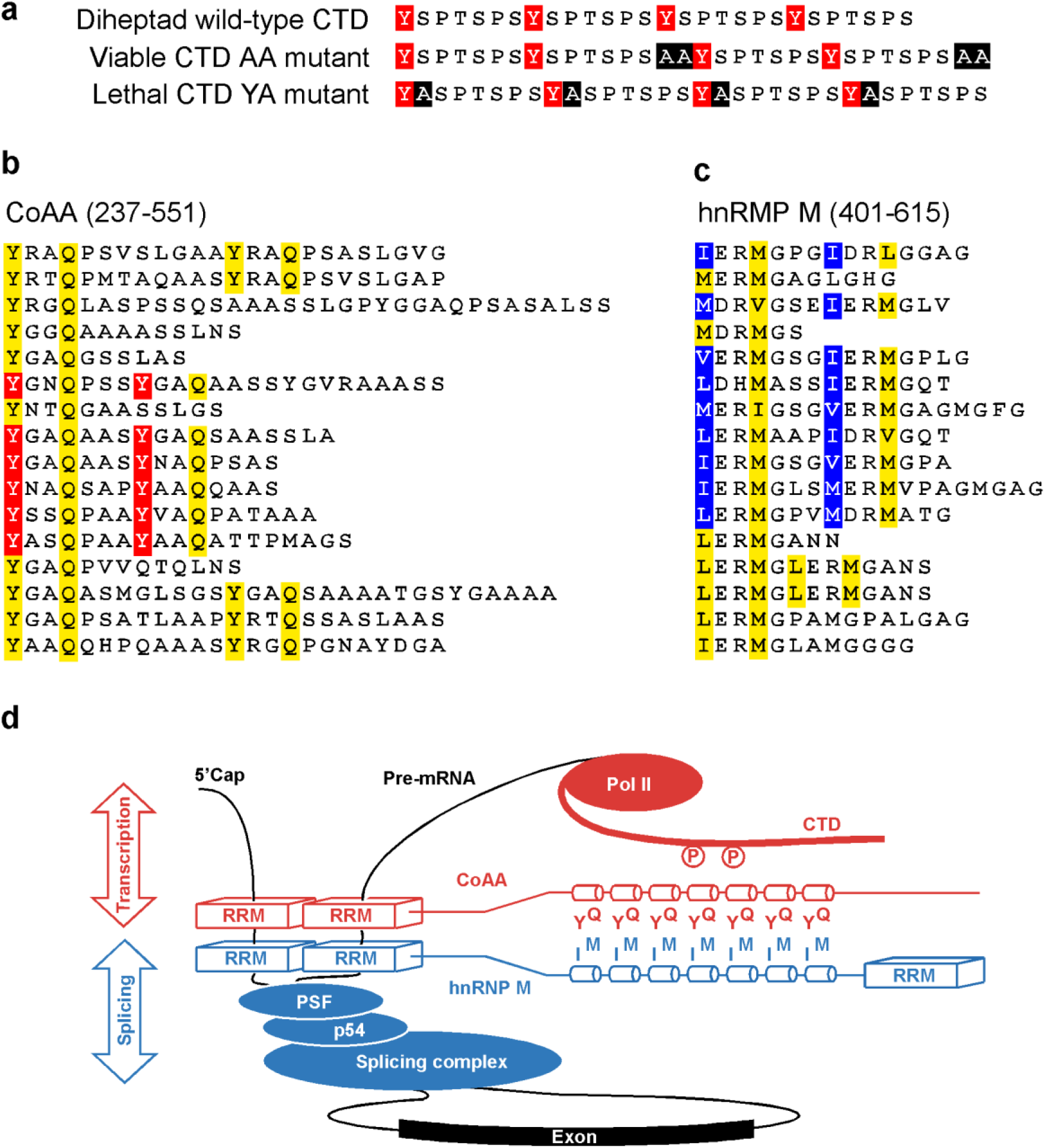
Pairs of heptad sequence repeats in the CTD of RNAP II, CoAA, and hnRNP M. **a**, Alignment of mutated CTD-YA and CTD-AA sequences with the wild-type CTD sequence. Alanine insertions are highlighted in black. Four heptad repeats are shown with tyrosine residues in red. **b**, CoAA 27-copy YxxQ repeats (237-551) are highlighted in yellow with tyrosines of heptad pairs in red. **c**, hnRNP M 27-copy IERM repeats (401-615) are highlighted in yellow with hydrophobic residues of heptad pairs in blue. Numbers indicate amino acids. **d**, Hypothetic model showing CoAA and hnRNP M in the interface of transcription and splicing regulations. Transcribed mRNA is shown in black. The coordination among transcription and splicing may be regulated by a myriad of factors including but not limited to CARM1.

**Supplementary Figure S5.**
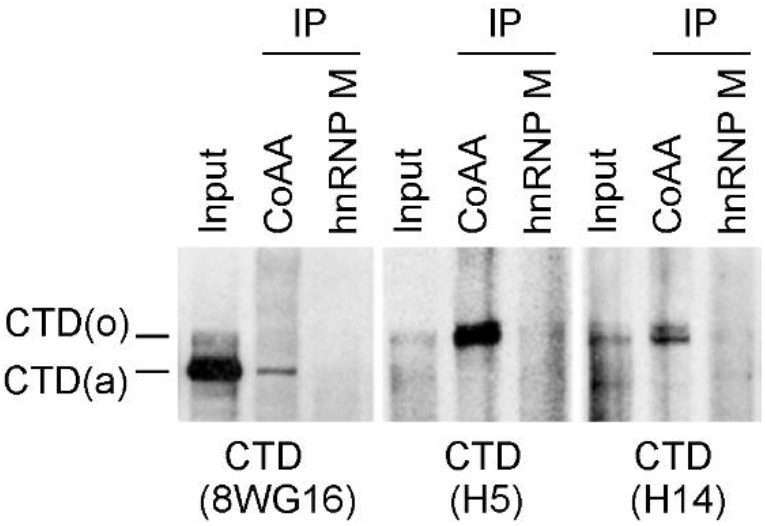
CoAA interacts with RNAP II. HeLa nuclear extracts were immunoprecipitated by anti-CoAA or anti-hnRNP M. Bound proteins were detected by Western blotting using anti-CTD (8WG16, H5, or H14). SDS-PAGE was 6% gel.

**Supplementary Figure S6.**
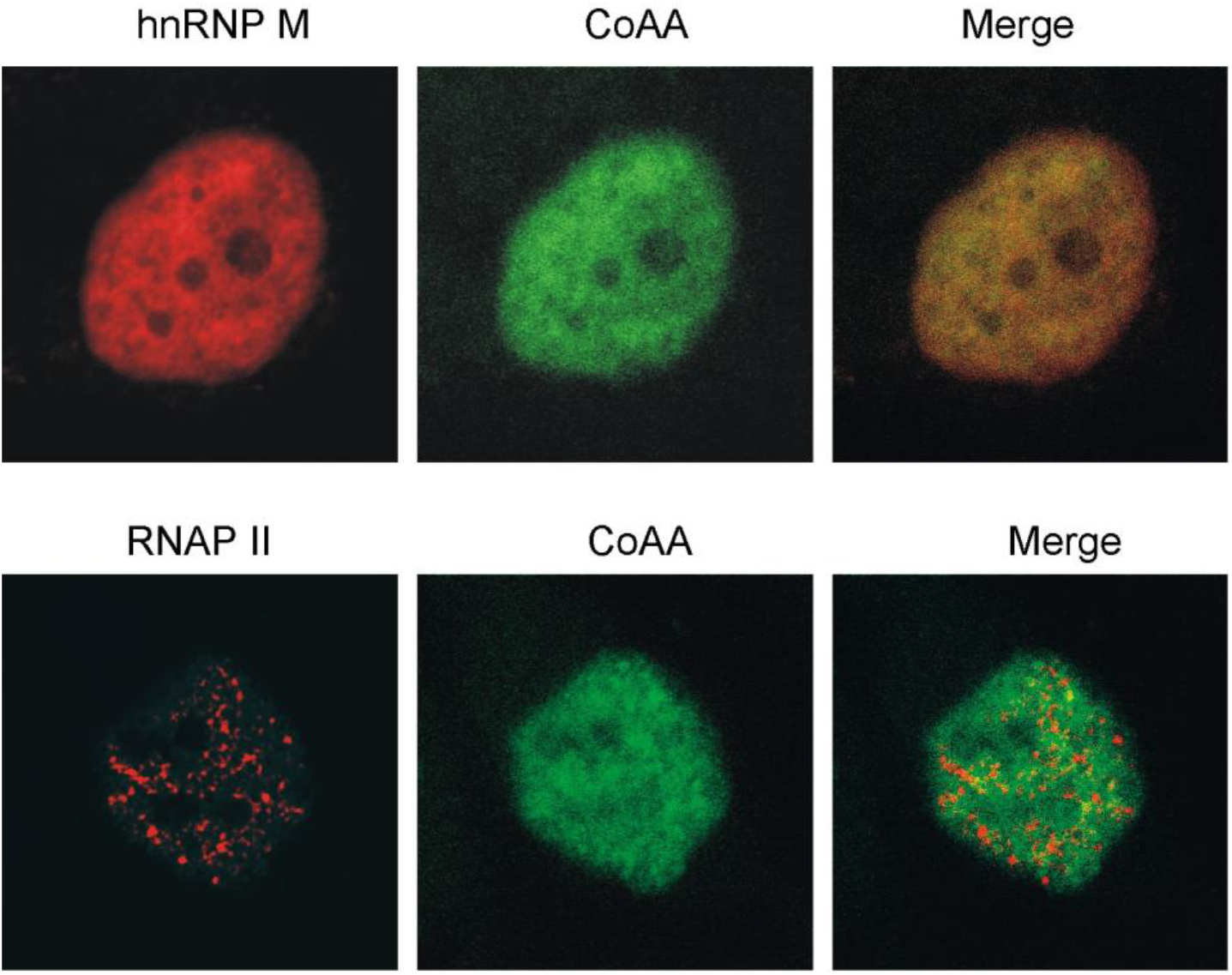
Nuclear localization of hnRNP M, CoAA, and RNAP II. HeLa cells were methanol fixed, and the nuclei were analyzed by immunofluorescent double staining using anti-CoAA, anti-hnRNP M or anti-CTD (8WG16) antibodies. Anti-rabbit FITC and anti-mouse Cy3 were secondary antibodies.

**Supplementary Figure S7.**
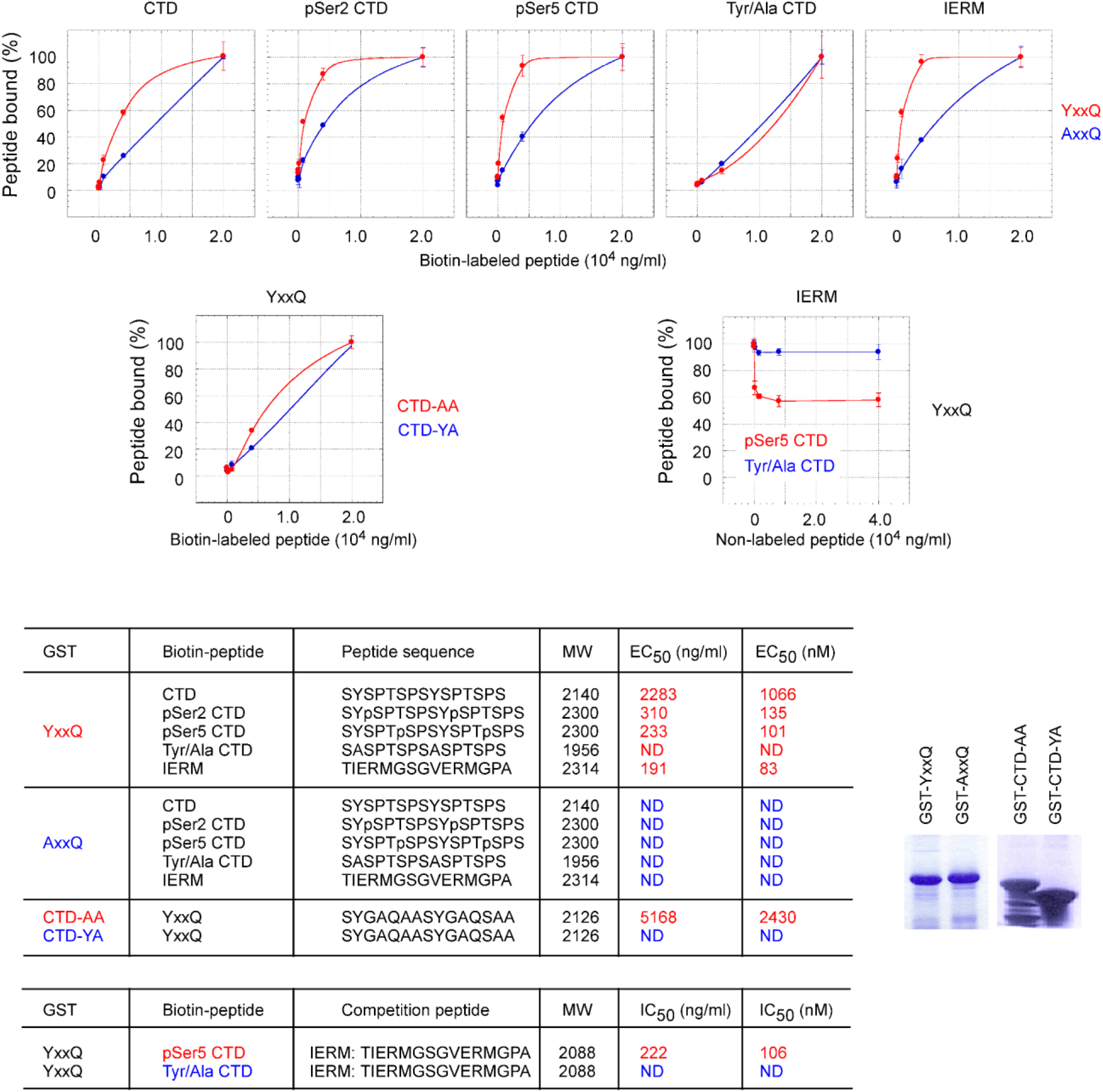
*In vitro* binding of diheptad peptides. **a-c**, The data sets in Fig. 2 were analyzed for the percentage of peptide bound as a function of peptide concentration. **d**, Diheptad peptide sequences, peptide molecular weight, and determined EC_50_ and IC_50_ values are listed with color corresponding to graphs. ND indicates the binding affinity below detectable level. In biotin-labeled peptides, two lysine residues and a C6 spacer (aminocaproic acid) are inserted between biotin and peptides in the format: [Biotin]-KK-C6-peptide. **e**, Coomassie blue staining of recombinant GST fusion proteins used in the assays.

**Supplementary Figure S8.**
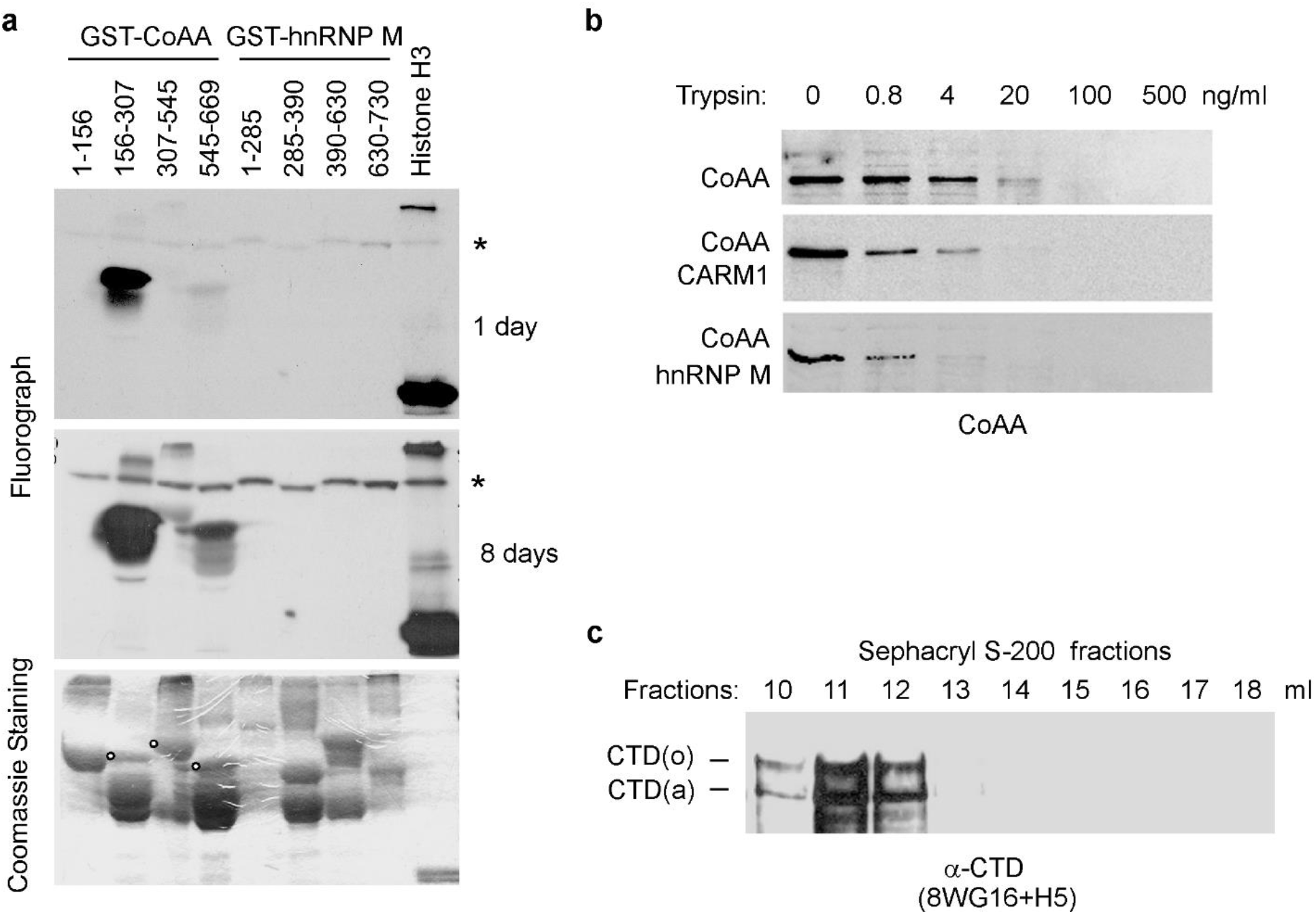
Arginine methylation of CoAA by CARM1. **a**, CoAA but not hnRNP M is methylated by CARM1. GST fusion proteins of CoAA or hnRNP M were tested for arginine methylation using CARM1 methyltransferase and labeled with [3H]-AdoMet *in vitro* using Histone H3 as a positive control. Auto-methylated CARM1 is indicated by asterisks. Same gel was fluorographed for methylation and Coomassie blue stained for protein loading. Exposure time was either 1 (shown in Fig. 4a) or 8 days as indicated. Positive bands were indicated by open circles. Degraded GST fragments were not methylated. **b**, Arginine methylation by CARM1 and hnRNP M interaction increase proteolytic sensitivity of CoAA. CoAA from the Mono Q fraction of 0.3 M NaCl (0.1 mg/ml) was either CARM1 methylated or incubated with His-tagged recombinant hnRNP M (0.02 mg/ml) before being subjected to proteolysis with increasing amounts of trypsin (1, 0.8, 4, 20, 100, 500 ng/ml). Proteolytic products were analyzed by anti-CoAA Western blot. **c**, Partial purification of RNAP II. HeLa nuclear extract was fractionated by gel filtration using a Sephacryl S-200 HR 10/30 column. Eluted fractions at 10-12 ml were collected for further Mono Q ion exchange purification. Each fraction was blotted with combined anti-CTD antibodies 8WG16 (1:200) and H5 (1:200).

## METHODS

### Plasmids and Antibodies

CoAA, MMTV-luciferase and glucocorticoid receptor plasmids are previously described (16,34). hnRNP M and hnRNP M1 were isolated by RT-PCR from HeLa cells and inserted into pcDNA3. CoAA and hnRNP M fragments were inserted in-frame into pM vector (Clontech) containing Gal4 DNA-binding domain to produce Gal4-fusion proteins. 5XGAL-luciferase reporter was from Stratagene. pETv5 was from Dr. Harald Konig (35). GST-CTD-YA and GST-CTD-AA were from Dr. John W. Stiller (13). pSG5-CARM1 was from Dr. Michael R. Stallcup. GST-CTD was from Dr. James L. Manley. The AxxQ and FxxQ mutants were generated by the substitution of 27 tyrosines with 27 alanines or phenylalanines using gene synthesis (MCLab) and verified by sequencing and Western blots. Rabbit polyclonal anti-CoAA was previously generated to against the RRM domains (1-156). Commercial antibodies are as follows: anti-FLAG M2 (F-3156, Sigma); anti-PSF (P-2860, Sigma); anti-Myc (46-0603, Invitrogen); anti-hnRNP M (2A6), anti-hnRNP M/M1 (1D8), (sc-20001, sc-20002, Santa Cruz); and anti-RNAP II CTD (8WG16, MMS-126R; H5, MMS-129R; H14, MMS-134R; Covance).

### Mass Spectrometry

GST-YxxQ (10 μg) was incubated with HeLa nuclear extracts in the binding buffer (20 mM Tris-HCl [pH 7.6], 50 mM NaCl, 75 mM KCl, 1 mM EDTA, 10% glycerol, 0.1% Triton X-100, 1 mM DTT and protease inhibitors). Bound proteins excised from preparative SDS-PAGE were washed with 50% acetonitrile and subjected to mass spectrometry analysis at the Harvard Microchemistry Facility. Sequence analysis included proteolytic digestion, microcapillary HPLC nano-electrospray tandem ion trap mass spectrometry, and MS/MS peptide sequence determination. Identifications were made with the Algorithm Sequest program.

### Recombinant Protein Binding Assays

*In vitro* binding assays were performed by incubating GST-YxxQ resin (20 μl, 2 μg) and ^35^S-methionine-labeled, *in vitro* translated hnRNP M protein fragments (5 μl) produced by rabbit reticulocyte lysate using TNT Quick Coupled Transcription/Translation Systems (Promega). Proteins were incubated at 4°C for 2 hours in the binding buffer above. Bound proteins were washed 3 times with the binding buffer and subjected to SDS-PAGE and autoradiography. For CTD phosphorylation *in vitro*, GST fusion proteins of the CTD and its mutants CTD-AA and CTD-YA were phosphorylated by Cdc2 kinase (NEB) using ^32^P-γ-ATP for 30 minutes at 30°C before incubated with immunoprecipitated CoAA or hnRNP M. Partially degraded GST-CTD protein fragments were not able to be phosphorylated but present in the system. Bound proteins were detected by autoradiography.

### Western Blotting and Co-immunoprecipitation (Co-IP)

Nuclear extracts were isolated by incubating cells in buffer A (20 mM HEPES, pH 7.4, 10 mM KCl, 1 mM EDTA, 1 mM EGTA, 0.1% Triton X-100, 1 mM DTT) with addition of leupeptin, aprotinin and trypsin inhibitor at 10 mg/ml for 15 min on ice. The cell pellets were then extracted in buffer B (20 mM HEPES, pH 7.4, 420 mM NaCl, 10 mM KCl, 1 mM EDTA, 1 mM EGTA, 0.5 mM MgCl2, 1 mM DTT) with protease inhibitors for 30 min. Nuclear extracts were further filtered using a 0.65 μm core size spin column (Millipore) to completely remove insoluble cellular debris. For coimmunoprecipitation, 5 ul of antibodies were captured by Protein A/G agarose (Santa Cruz) for IgG or by Immobilized Protein L agarose (Pierce) for IgM. The immune complexes were washed before incubating with 1:10 diluted nuclear extracts in the binding buffer. The precipitates were washed and subjected to Western blotting analysis using appropriate primary antibodies. The blots were detected with the ECL system (Amersham Pharmacia).

### Peptide Binding Assays

White Microlite 2+ 96-well plates were coated overnight at 37°C with 500 ng/well of either GST-YxxQ protein (307-545) or GST-CTD-AA or GST-CTD-YA(13) in the presence of 5 μg/well of BSA. The coated plates were blocked with 5% BSA in 0.01 M Tris-HCl/0.15 M NaCl for 1 hour. The wells were then incubated overnight in duplicates with biotin-labeled CTD, IERM or YxxQ peptides at increasing concentrations of 6.4, 32, 160, 800, 4000, and 20000 ng/ml. Competition assays were performed using increasing concentrations of nonlabeled IERM peptide at 12.8, 64, 320, 1600, 8000, 40000 ng/ml and a constant concentration of biotin-labeled CTD peptides at 10^4^ ng/ml. The plates were quickly washed 4 times with the binding buffer (20 mM HEPES, pH 7.4, 50 mM NaCl, 75 mM KCl, 1 mM EDTA, 0.05% Triton X-100, 10% glycerol, 1 mM DTT), and dried again overnight to prevent disruption of the binding equilibrium during later washing steps. Bound biotin peptides were then detected by incubating with 1 U/ml of HRP-conjugated streptavidin (Roche) on ice for 45 min in PBS/0.1% Triton X-100. Washed plates were added with 50 μL/well ECL detecting reagents and read on a Dynex luminometer for 3 seconds. EC_50_ and IC_50_ values were determined with sigmoidal non-linear progression curve fit in Schild plots using KaleidaGraph program. Peptide sequences: CTD, SYSPTSPSYSPTSPS; pSer2 CTD, SYpSPTSPSYpSPTSPS; pSer5 CTD, SYSPTpSPSYSPTpSPS; Tyr/Ala CTD, SASPTSPSASPTSPS; IERM, TIERMGSGVERMGPA; YxxQ, SYGAQAASYGAQSAA. In biotin-labeled peptides, two lysine residues and a C6 spacer (aminocaproic acid) are inserted between biotin and peptides in the format: [Biotin]-KK-C6-peptide.

### Cell Culture and Transient Transfection

CV1 cells were maintained in DMEM supplemented with 10% fetal bovine serum and 5 μg/μl penicillin/streptomycin in 5% CO2 at 37 °C. In GAL4 reporter assays, cells in 24-well plates were transfected in triplicates with the 5XGAL-luciferase reporter (100 ng) and GAL4 fusion plasmids (200 ng) using Lipofectin (Life Technologies, Inc.). In MMTV reporter assays, cells were transfected with MMTV-luciferase (100 ng), glucocorticoid receptor (10 ng), CoAA (200 ng), or hnRNP M (200 ng) plasmids per well and induced by dexamethasone (Dex) (100 nM) for an additional 16 hours after transfection. Luciferase activities were measured by a Dynex luminometer. Data are shown as means of triplicate transfections ± standard errors.

### Alternative Splicing Analysis

The pETv5 alternative splicing reporter contains the CD44 variable exon 5 and its adjacent intron sequences inserted between pre-proinsulin exons 2 and exon 3(36). The detecting primers on pre-proinsulin exons distinguish the minigene from the endogenous transcripts. 293 cells were transiently transfected with pETv5 minigene driven by RSV promoter together with CoAA or hnRNP M expression vectors or with their siRNA (100 nM) as indicated. Total RNA from cells was prepared with Trizol reagent (Invitrogen), treated with DNase I, and followed by RT-PCR. RT-PCR primers are as follows: sense, AGTGGATCCGCTTCCTGCCCC; antisense, CTGCCGGGCCACCTCCAGTGCC. The target sequence of siRNA (Dharmacon) is 5’-AGAUUAUCCAUGCAUUACA-3’ for hnRNP M, 5’-GUAACCAGCCAUCCUCUUA-3’ for CoAA, and 5’-UAGCGACUAAACACAUCAA-3’ for the control.

### Chromatin Immunoprecipitation (ChIP)

HeLa cells were incubated with 1% formaldehyde for 10 min to crosslink proteins and DNA, and the reaction was stopped by 125 mM glycine. Cells were lysed and sonicated in buffer containing 20 mM Tris pH 8.0, 75 mM NaCl, 75 mM KCl, 1 mM EDTA, 1 mM EGTA, 1% Triton X-100, 10% glycerol, 1 mM DTT, and protease inhibitors. Immunoprecipitation was carried out using salmon sperm DNA-blocked protein A/G resin (Upstate) and individual antibodies. The crosslinking was reversed by eluting with 0.1 M NaHCO3, 1% SDS, 0.3 M NaCl at 65°C for 4 hours. Purified DNA (Qiagen kit) was subjected to real-time PCR analysis. Primer pairs used on the CoAA gene are listed in Supplementary Table S2.

### Protein Chromatography

RNAP II was partially purified from HeLa nuclear extracts using Sephacryl S-200 HR 10/30 gel filtration column with buffer A (20 mM Tris pH 8.0, 1 mM EDTA, and 1 mM DTT) containing 150 mM NaCl (Supplementary Fig. S8c). CoAA or desalted RNAP II fractions were further purified by a Mono Q column (5 ml) equilibrated with buffer A containing 10 mM NaCl and step eluted with buffer A containing 0.1-0.8 M NaCl. Fractions were analyzed by Western blotting using anti-CTD 8WG16 (1:200), H5 (1:200), CoAA (1:200) or hnRNP M (1:500) antibodies. Mono Q 0.4-0.5 M NaCl fractions containing RNAP II were concentrated before use. CoAA was purified using Mono Q column only.

### Arginine Methylation by CARM1

GST fusion proteins of CoAA and hnRNP M fragments were *in vitro* arginine methylated by baculovirus expressed CARM1 (0.1 μg) in a 20 μl methylation reaction (20 mM Tris pH 8.0, 1 mM EDTA, and 200 mM NaCl) containing 1 μCi S-adenosyl-L-[methyl-^3^H] methionine (AdoMet, Amersham Bioscience) at 30°C for 60 min(37). The SDS gel containing labeled protein was stained with Coomassie blue, soaked with Amplify solution for 15 min, before fluorography (Supplementary Fig. S8a). In methylation-induced binding assays, CoAA from partially purified fractions was immunoprecipitated and washed with the binding buffer for three times before methylated by CARM1. Methylation of immunoprecipitated CoAA was performed under the same methylation conditions as above except using cold AdoMet (0.25 mM, Sigma) for 20 min. Methylated CoAA captured on beads was further incubated with purified RNAP II and analyzed by coimmunoprecipitation.

### Sequence Repeat Analysis

The uniqueness of the YxxQ repeats of CoAA and the IERM repeats of hnRNP M in the human genome was analyzed by ScanProsite at ExPASy Proteomics Server (http://ca.expasy.org), using protein databases at Swiss-Prot (release 44.3; 156998 entries) and TrEMBL (release 27.3; 1379120 entries). The diheptad sequence patterns in CoAA and hnRNP M were revealed by RADAR program (Rapid Automatic Detection and Alignment of Repeats www.ebi.ac.uk/Tools/Radar).

## References

1. Meinhart, A., Kamenski, T., Hoeppner, S., Baumli, S., and Cramer, P. (2005) A structural perspective of CTD function. Genes Dev 19, 1401–1415

2. Orphanides, G., and Reinberg, D. (2002) A unified theory of gene expression. Cell 108, 439–451

3. Fong, N., Bird, G., Vigneron, M., and Bentley, D. L. (2003) A 10 residue motif at the C-terminus of the RNA pol II CTD is required for transcription, splicing and 3’ end processing. Embo J 22, 4274–4282

4. Kaneko, S., and Manley, J. L. (2005) The mammalian RNA polymerase II C-terminal domain interacts with RNA to suppress transcription-coupled 3’ end formation. Mol Cell 20, 91–103

5. Cho, E. J., Kobor, M. S., Kim, M., Greenblatt, J., and Buratowski, S. (2001) Opposing effects of Ctk1 kinase and Fcp1 phosphatase at Ser 2 of the RNA polymerase II C-terminal domain. Genes Dev 15, 3319–3329

6. Corden, J. L., and Patturajan, M. (1997) A CTD function linking transcription to splicing. Trends Biochem Sci 22, 413–416

7. Xu, Y. X., Hirose, Y., Zhou, X. Z., Lu, K. P., and Manley, J. L. (2003) Pin1 modulates the structure and function of human RNA polymerase II. Genes Dev 17, 2765–2776

8. Chapman, R. D., Heidemann, M., Albert, T. K., Mailhammer, R., Flatley, A., Meisterernst, M., Kremmer, E., and Eick, D. (2007) Transcribing RNA polymerase II is phosphorylated at CTD residue serine-7. Science 318, 1780–1782

9. Yurko, N. M., and Manley, J. L. (2018) The RNA polymerase II CTD “orphan” residues: Emerging insights into the functions of Tyr-1, Thr-4, and Ser-7. Transcription 9, 30–40

10. Shah, N., Maqbool, M. A., Yahia, Y., El Aabidine, A. Z., Esnault, C., Forne, I., Decker, T. M., Martin, D., Schuller, R., Krebs, S., Blum, H., Imhof, A., Eick, D., and Andrau, J. C. (2018) Tyrosine-1 of RNA Polymerase II CTD Controls Global Termination of Gene Transcription in Mammals. Molecular cell 69, 48–61 e46

11. Schuller, R., Forne, I., Straub, T., Schreieck, A., Texier, Y., Shah, N., Decker, T. M., Cramer, P., Imhof, A., and Eick, D. (2016) Heptad-Specific Phosphorylation of RNA Polymerase II CTD. Molecular cell 61, 305–314

12. Jeronimo, C., Collin, P., and Robert, F. (2016) The RNA Polymerase II CTD: The Increasing Complexity of a Low-Complexity Protein Domain. Journal of molecular biology 428, 2607–2622

13. Liu, P., Greenleaf, A. L., and Stiller, J. W. (2008) The Essential Sequence Elements Required for RNAP II C-terminal Domain Function in Yeast and their Evolutionary Conservation. Mol Biol Evol 25, 719–727

14. Stiller, J. W., and Cook, M. S. (2004) Functional unit of the RNA polymerase II C-terminal domain lies within heptapeptide pairs. Eukaryot Cell 3, 735–740

15. McKenna, N. J., Lanz, R. B., and O’Malley, B. W. (1999) Nuclear receptor coregulators: cellular and molecular biology. Endocr Rev 20, 321–344

16. Iwasaki, T., Chin, W. W., and Ko, L. (2001) Identification and characterization of RRM-containing coactivator activator (CoAA) as TRBP-interacting protein, and its splice variant as a coactivator modulator (CoAM). J Biol Chem 276, 33375–33383

17. Auboeuf, D., Dowhan, D. H., Li, X., Larkin, K., Ko, L., Berget, S. M., and O’Malley, B. W. (2004) CoAA, a nuclear receptor coactivator protein at the interface of transcriptional coactivation and RNA splicing. Mol Cell Biol 24, 442–453

18. Sui, Y., Yang, Z., Xiong, S., Zhang, L., Blanchard, K. L., Peiper, S. C., Dynan, W. S., Tuan, D., and Ko, L. (2007) Gene amplification and associated loss of 5’ regulatory sequences of *CoAA* in human cancers. Oncogene 26, 822–835

19. Yang, Z., Sui, Y., Xiong, S., Liour, S. S., Phillips, A. C., and Ko, L. (2007) Switched Alternative Splicing of Oncogene CoAA during embryonal carcinoma stem cell differentiation. Nucleic Acids Res 35, 1919–1932

20. Kang, Y. K., Schiff, R., Ko, L., Wang, T., Tsai, S. Y., Tsai, M. J., and O’Malley, B. W. (2008) Dual roles for CoAA and its counterbalancing isoform CoAM in human kidney cell tumorigenesis. Cancer Research In press

21. Ko, L. (2019) Human solid cancer decoded. Zenodo preprint, http://doi.org/10.5281/zenodo.3236836

22. Perani, M., Antonson, P., Hamoudi, R., Ingram, C. J., Cooper, C. S., Garrett, M. D., and Goodwin, G. H. (2005) The proto-oncoprotein SYT interacts with SYT-interacting protein/co-activator activator (SIP/CoAA), a human nuclear receptor co-activator with similarity to EWS and TLS/FUS family of proteins. J Biol Chem 280, 42863–42876

23. Kim, J., and Pelletier, J. (1999) Molecular genetics of chromosome translocations involving EWS and related family members. Physiol Genomics 1, 127–138

24. Datar, K. V., Dreyfuss, G., and Swanson, M. S. (1993) The human hnRNP M proteins: identification of a methionine/arginine-rich repeat motif in ribonucleoproteins. Nucleic Acids Res 21, 439–446

25. Gattoni, R., Mahe, D., Mahl, P., Fischer, N., Mattei, M. G., Stevenin, J., and Fuchs, J. P. (1996) The human hnRNP-M proteins: structure and relation with early heat shock-induced splicing arrest and chromosome mapping. Nucleic Acids Res 24, 2535–2542

26. Kafasla, P., Patrinou-Georgoula, M., Lewis, J. D., and Guialis, A. (2002) Association of the 72/74-kDa proteins, members of the heterogeneous nuclear ribonucleoprotein M group, with the pre-mRNA at early stages of spliceosome assembly. Biochem J 363, 793–799

27. Kornblihtt, A. R. (2005) Promoter usage and alternative splicing. Curr Opin Cell Biol 17, 262–268

28. Bedford, M. T. (2007) Arginine methylation at a glance. J Cell Sci 120, 4243–4246

29. Cheng, D., Cote, J., Shaaban, S., and Bedford, M. T. (2007) The arginine methyltransferase CARM1 regulates the coupling of transcription and mRNA processing. Mol Cell 25, 71–83

30. Liu, Q., and Dreyfuss, G. (1995) In vivo and in vitro arginine methylation of RNA-binding proteins. Mol Cell Biol 15, 2800–2808

31. Belyanskaya, L. L., Gehrig, P. M., and Gehring, H. (2001) Exposure on cell surface and extensive arginine methylation of ewing sarcoma (EWS) protein. J Biol Chem 276, 18681–18687

32. Meinhart, A., and Cramer, P. (2004) Recognition of RNA polymerase II carboxyterminal domain by 3’-RNA-processing factors. Nature 430, 223–226

33. Brooks, Y. S., Wang, G., Yang, Z., Smith, K. K., Bieberich, E., and Ko, L. (2009) Functional pre-mRNA trans-splicing of coactivator CoAA and corepressor RBM4 during stem/progenitor cell differentiation. J Biol Chem 284, 18033–18046

34. Ko, L., Cardona, G. R., and Chin, W. W. (2000) Thyroid hormone receptor-binding protein, an LXXLL motif-containing protein, functions as a general coactivator. Proc Natl Acad Sci U S A 97, 6212–6217

35. Konig, H., Ponta, H., and Herrlich, P. (1998) Coupling of signal transduction to alternative pre-mRNA splicing by a composite splice regulator. Embo J 17, 2904–2913

36. Konig, H., Moll, J., Ponta, H., and Herrlich, P. (1996) Trans-acting factors regulate the expression of CD44 splice variants. Embo J 15, 4030–4039

37. Xu, W., Chen, H., Du, K., Asahara, H., Tini, M., Emerson, B. M., Montminy, M., and Evans, R. M. (2001) A transcriptional switch mediated by cofactor methylation. Science 294, 2507–2511

