## Supplementary Tables S1-S2 for "Oncoprotein CoAA repeats interact with RNA polymerase II CTD repeats"

### Supplementary Table S1-S2

### Oncoprotein CoAA repeats interact with RNA polymerase II CTD repeats

Shiqin Xiong,[1] Yang S. Brooks,[1] Zheqiong Yang,[1] Jiacai Wu,[2] Liyong Zhang,[1] William S. Dynan,[1] Wei Xu,[2] Bert W. O'Malley,[3] and Lan Ko [1*]

### Supplementary Table S1

**Ion Current (Sum):** 2.0e7
**Ion Current (Avg):** 6.2e5
**# MS/MS Spectra:** 415

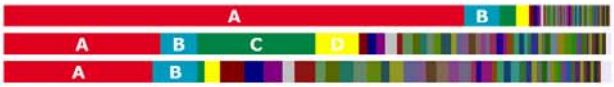


**A** gi|14141152 **MS/MS Spectra:** 102 **Sum TIC:** 1.6e7 **Avg TIC:** 1.6e5
heterogeneous nuclear ribonucleoprotein M isoform a; heterogenous nuclear ribonucleoprotein M4; N-acetylglucosamine receptor 1; M4 protein; heterogenous nuclear ribonucleoprotein M [Homo sapiens] gi|13111793|gb|AAH00138.2| Heterogeneous nuclear ribonucleoprotein M, isof

| | Sequence | Reference | TIC | Ions | Scan |
|---|---|---|---|---|---|
| (-) | VGEVTYVELLM*DAEGK | gi\|14141152 +9 | 5.8e4 | 24/30 | 5084-6010 |
| (-) | VGEVTYVELLMDAEGK | gi\|14141152 +9 | 6.0e3 | 22/30 | 5992 |
| (-) | PQQLPHGLGGIGM*GLGPGGQPIDANHLNK | gi\|14141152 +9 | 7.9e3 | 41/112 | 4937 |
| (-) | MGPLGLDHMASSIER | gi\|14141152 +14 | 7.0e4 | 22/28 | 4523-4935 |
| (-) | MGLAMGGGGGASFDR | gi\|14141152 +8 | 5.7e4 | 24/28 | 3994-4367 |
| (-) | GIGMGNIGPAGMGMEGIGFGINK | gi\|14141152 +8 | 4.8e3 | 26/44 | 5852 |
| (-) | LKEVFSMAGVVVR | gi\|14141152 +11 | 4.3e3 | 21/24 | 5094 |
| (-) | GIGMGNIGPAGMGM*EGIGFGINK | gi\|14141152 +8 | 2.2e4 | 40/88 | 5497-5690 |
| (-) | M*EEESGAPGVPSGNGAPGPK | gi\|14141152 +4 | 1.1e5 | 25/38 | 3350-3478 |
| (-) | LGSTVFVANLDYK | gi\|14141152 +11 | 1.3e6 | 19/24 | 4962-5843 |
| (-) | MGPAM*GPALGAGIER | gi\|14141152 +13 | 8.6e5 | 21/28 | 4042-4502 |
| (-) | GIGM*GNIGPAGMGM*EGIGFGINK | gi\|14141152 +8 | 6.2e4 | 37/88 | 5104-5638 |
| (-) | LGSTVFVANLDYK | gi\|14141152 +11 | 8.2e3 | 25/48 | 4982 |
| (-) | INEILSNALK | gi\|14141152 +13 | 1.4e6 | 16/18 | 4435-6478 |
| (-) | GIGMGNIGPAGM*GM*EGIGFGINK | gi\|14141152 +8 | 1.2e5 | 25/44 | 5132-5569 |
| (-) | AFITNIPFDVK | gi\|14141152 +12 | 8.8e5 | 17/20 | 5399-12514 |
| (-) | EKVGEVTYVELLM*DAEGK | gi\|14141152 +9 | 2.1e3 | 19/34 | 5555 |
| (-) | EVFSMAGVVVR | gi\|14141152 +11 | 1.7e5 | 16/20 | 4972 |
| (-) | GNFGGSFAGSFGGAGGHAPGVAR | gi\|14141152 +13 | 1.3e4 | 20/44 | 4594 |
| (-) | ADILEDKDGK | gi\|14141152 +11 | 1.9e5 | 16/18 | 2800-3187 |
| (-) | MEEESGAPGVPSGNGAPGPK | gi\|14141152 +4 | 3.2e5 | 24/38 | 3375-3518 |
| (-) | MGGMEGPFGGGMENMGR | gi\|14141152 +8 | 3.3e3 | 19/32 | 4822 |
| (-) | GIGMGNIGPAGM*GMEGIGFGINK | gi\|14141152 +8 | 3.1e4 | 23/44 | 5474-5733 |
| (-) | GEGERPAQNEK | gi\|14141152 +6 | 1.1e4 | 16/20 | 1422 |
| (-) | M*GLAM*GGGGGASFDR | gi\|14141152 +8 | 1.6e5 | 21/28 | 3423-4215 |
| (-) | M*GAGM*GFGLER | gi\|14141152 +14 | 2.9e5 | 16/20 | 3702-4532 |
| (-) | M*GPAM*GPALGAGIER | gi\|14141152 +13 | 2.5e5 | 18/28 | 3894-4413 |
| (-) | GGNRFEPYANPTK | gi\|14141152 +8 | 1.5e4 | 16/24 | 3525 |
| (-) | VKEDPDGEHAR | gi\|14141152 +8 | 4.3e4 | 17/20 | 1420-2007 |
| (-) | M*GLAMGGGGGASFDR | gi\|14141152 +8 | 8.3e3 | 18/28 | 4223-12505 |
| (-) | MGANSLER | gi\|14141152 +27 | 2.3e5 | 13/14 | 2000-3052 |
| (-) | INEILSNALKR | gi\|14141152 +13 | 1.5e4 | 16/20 | 4229-4445 |
| (-) | M*GLAMGGGGGASFDR | gi\|14141152 +8 | 1.3e5 | 19/28 | 3449-4500 |
| (-) | YRAFITNIPFDVK | gi\|14141152 +10 | 2.0e3 | 17/24 | 5609 |
| (-) | FNECGHVLYADIK | gi\|14141152 +13 | 6.0e4 | 24/48 | 4332-4464 |
| (-) | MGLVMDR | gi\|14141152 +13 | 6.2e5 | 11/12 | 168-9674 |
| (-) | M*GAGLGHGMDR | gi\|14141152 +13 | 1.5e5 | 16/20 | 2095-3263 |
| (-) | QGGGGGGGSVPGIER | gi\|14141152 +8 | 1.2e6 | 16/28 | 3327 |
| (-) | MGPLGLDHM*ASSIER | gi\|14141152 +14 | 2.3e5 | 28/56 | 4055-12557 |
| (-) | GCAVVEFK | gi\|14141152 +10 | 1.5e5 | 12/14 | 3664-3732 |
| (-) | KACQIFVR | gi\|14141152 +14 | 1.4e5 | 12/14 | 3492-3507 |
| (-) | LGGAGMER | gi\|14141152 +8 | 6.0e5 | 12/14 | 79-12515 |
| (-) | M*GAGLGHGM*DR | gi\|14141152 +13 | 4.2e4 | 15/20 | 2038-3040 |
| (-) | M*GPLGLDHM*ASSIER | gi\|14141152 +14 | 1.8e5 | 19/28 | 3977-4595 |
| (-) | LGSTVFVANLDYK | gi\|14141152 +11 | 3.4e3 | 15/24 | 5222 |
| (-) | VGQTIER | gi\|14141152 +14 | 4.7e5 | 11/12 | 793-12504 |
| (-) | MGANNLER | gi\|14141152 +14 | 1.5e5 | 12/14 | 1592-2982 |
| (-) | MGPLGLDHMASSIER | gi\|14141152 +14 | 2.3e5 | 28/56 | 4507-9085 |
| (-) | GGNRFEPYANPTKR | gi\|14141152 +8 | 2.0e4 | 15/26 | 3500 |
| (-) | MGPAM*GPALGAGIER | gi\|14141152 +13 | 2.2e4 | 19/28 | 4433-12541 |
| (-) | M*GGMEGPFGGGMENMGR | gi\|14141152 +8 | 2.5e4 | 29/64 | 4267-4527 |
| (-) | M*GANNLER | gi\|14141152 +14 | 1.9e5 | 12/14 | 912-12694 |
| (-) | VKEDPDGEHAR | gi\|14141152 +8 | 8.0e4 | 21/40 | 1482 |
| (-) | DKFNECGHVLYADIK | gi\|14141152 +13 | 3.5e5 | 24/56 | 4342 |
| (-) | VKEDPDGEHAR | gi\|14141152 +8 | 1.2e5 | 24/40 | 427-9745 |
| (-) | M*GPLGLDHMASSIER | gi\|14141152 +14 | 3.3e5 | 17/28 | 4040-4910 |
| (-) | MGPLGLDHM*ASSIER | gi\|14141152 +14 | 1.6e5 | 27/56 | 4037-4947 |
| (-) | ADILEDKDGK | gi\|14141152 +11 | 2.1e3 | 12/18 | 3104 |
| (-) | M*VPAGM*GAGLER | gi\|14141152 +9 | 2.3e5 | 17/22 | 3230-3843 |
| (-) | LKEVFSMAGVVVR | gi\|14141152 +11 | 1.3e4 | 23/48 | 5087-8919 |
| (-) | GNFGGSFAGSFGGAGGHAPGVAR | gi\|14141152 +13 | 4.0e5 | 27/88 | 4652-7260 |
| (-) | ACQIFVR | gi\|14141152 +14 | 1.3e4 | 11/12 | 3677 |
| (-) | NLPFDFTWK | gi\|14141152 +14 | 8.3e4 | 11/16 | 5827-12499 |
| (-) | EKVGEVTYVELLM*DAEGK | gi\|14141152 +9 | 2.2e3 | 23/68 | 5525 |
| (-) | ADILEDKDGKSR | gi\|14141152 +11 | 5.7e3 | 14/22 | 2958 |
| (-) | MGGM*EGPFGGGMENMGR | gi\|14141152 +8 | 1.5e5 | 16/32 | 4154-4615 |
| (-) | M*GANSLER | gi\|14141152 +27 | 3.8e5 | 10/14 | 1605-9517 |
| (-) | M*GAGLGHGMDR | gi\|14141152 +13 | 4.1e4 | 15/20 | 1634-3132 |
| (-) | MEESMK | gi\|14141152 +18 | 1.8e4 | 8/10 | 1589-2037 |

| | | | | | |
|---|---|---|---|---|---|
| (-) | MGPVMDR | gi\|14141152 +14 | 8.5e4 | 10/12 | 3202 |
| (-) | AFITNIPFDVK | gi\|14141152 +12 | 7.7e3 | 13/20 | 5507-5513 |
| (-) | GCAVVEFK | gi\|14141152 +10 | 2.2e4 | 10/14 | 3684-3703 |
| (-) | SRGCAVVEFK | gi\|14141152 +10 | 1.2e4 | 12/18 | 3547 |
| (-) | LGGAGM*ER | gi\|14141152 +8 | 1.8e5 | 10/14 | 37-12759 |
| (-) | M*GLAMGGGGGASFDR | gi\|14141152 +8 | 6.7e3 | 16/28 | 4663 |
| (-) | AAEVLNK | gi\|14141152 +10 | 8.0e4 | 8/12 | 1804-2347 |
| (-) | MGLAMGGGGGASFDR | gi\|14141152 +8 | 7.3e2 | 13/28 | 5883 |
| (-) | NLPFDFTWK | gi\|14141152 +14 | 1.6e4 | 10/16 | 5835-5858 |
| (-) | MAAPIDR | gi\|14141152 +15 | 2.0e4 | 9/12 | 3148-3159 |
| (-) | KLKEVFSM*AGVVVR | gi\|14141152 +11 | 1.4e4 | 18/52 | 4438 |
| (-) | MGQTM*ER | gi\|14141152 +14 | 1.0e5 | 10/12 | 84-9882 |
| (-) | M*GAGMGFGLER | gi\|14141152 +14 | 6.4e5 | 14/20 | 3729-4454 |
| (-) | FGSGMNMGR | gi\|14141152 +14 | 1.2e4 | 11/16 | 3415-3447 |
| (-) | MGLSMER | gi\|14141152 +6 | 2.0e4 | 9/12 | 3594-3613 |
| (-) | LGSTVFVANLDYK | gi\|14141152 +11 | 7.1e2 | 11/24 | 4967 |
| (-) | MGGM*EGPFGGGM*ENMGR | gi\|14141152 +8 | 3.8e5 | 15/32 | 3907-4432 |
| (-) | M*GPAMGPALGAGIER | gi\|14141152 +13 | 3.6e3 | 13/28 | 4795 |
| (-) | INEILSNALK | gi\|14141152 +13 | 2.3e4 | 11/18 | 4547-4570 |
| (-) | EVFSMAGVVVR | gi\|14141152 +11 | 1.5e3 | 11/20 | 4994 |
| (-) | MGQTMER | gi\|14141152 +14 | 2.1e4 | 8/12 | 2052-2195 |
| (-) | M*GPLGLDHM*ASSIER | gi\|14141152 +14 | 7.8e5 | 22/56 | 3967-6344 |
| (-) | FGSGM*NMGR | gi\|14141152 +14 | 2.3e5 | 8/16 | 2804-3452 |
| (-) | FESPEVAER | gi\|14141152 +15 | 8.2e3 | 8/16 | 3437 |
| (-) | ACQIFVR | gi\|14141152 +14 | 3.2e3 | 7/12 | 3685 |
| (-) | VGQTIER | gi\|14141152 +14 | 4.6e4 | 7/12 | 2137-2318 |
| (-) | FEPYANPTK | gi\|14141152 +8 | 7.6e4 | 10/16 | 3448-3460 |
| (-) | MGPVMDR | gi\|14141152 +14 | 1.6e4 | 7/12 | 3225 |
| (-) | M*GLAMGGGGGASFDR | gi\|14141152 +8 | 1.5e3 | 11/28 | 5174 |
| (-) | AFITNIPFDVK | gi\|14141152 +12 | 5.4e2 | 10/20 | 8950 |
| (-) | FGSGMNMGR | gi\|14141152 +14 | 9.5e3 | 8/16 | 3475 |
| (-) | M*ATGLER | gi\|14141152 +14 | 4.5e3 | 6/12 | 2105 |
| (-) | GIGM*GNIGPAGMGM*EGIGFGINK | gi\|14141152 +8 | 5.1e3 | 12/44 | 5537 |

**B** gi|15022507 **MS/MS Spectra:** 31 **Sum TIC:** 1.2e6 **Avg TIC:** 3.7e4
coactivator activator [Homo sapiens]

| **Sequence** | | **Reference** | **TIC** | **Ions** | **Scan** |
|---|---|---|---|---|---|
| (-) | TQSSASLAASYAAQQHPQAAASYR | gi\|15022507 +5 | 7.3e3 | 24/46 | 4008 |
| (-) | IFVGNVSAACTSQELR | gi\|15022507 +7 | 4.6e3 | 22/30 | 4440 |
| (-) | ASYVAPLTAQPATYR | gi\|15022507 +5 | 1.6e5 | 21/28 | 4317-4327 |
| (-) | AQPSVSLGAAYR | gi\|15022507 +5 | 2.0e5 | 20/22 | 3940-3972 |
| (-) | AQPSVSLGAPYR | gi\|15022507 +5 | 1.8e4 | 18/22 | 3984 |
| (-) | LPDAHSDYAR | gi\|15022507 +3 | 3.6e4 | 16/18 | 3177 |
| (-) | IFVGNVSAACTSQELR | gi\|15022507 +7 | 1.5e4 | 29/60 | 4452 |
| (-) | KGPGLAVQSGDK | gi\|15022507 +1 | 2.3e4 | 17/22 | 3017-3134 |
| (-) | LAELSDYR | gi\|15022507 +3 | 1.1e5 | 13/14 | 3777-3783 |
| (-) | AAQMHSGYQR | gi\|15022507 +3 | 1.5e4 | 14/18 | 2093 |
| (-) | RLPDAHSDYAR | gi\|15022507 +3 | 4.5e4 | 15/20 | 3137-3164 |
| (-) | TQPM*TAQAASYR | gi\|15022507 +1 | 2.0e4 | 16/22 | 2948-3208 |
| (-) | YSGSYNDYLR | gi\|15022507 +3 | 1.2e5 | 14/18 | 3715-3947 |
| (-) | GPGLAVQSGDK | gi\|15022507 +1 | 6.1e3 | 16/20 | 3069-3075 |
| (-) | AIEALHGHELRPGR | gi\|15022507 +6 | 2.2e4 | 26/52 | 3578 |
| (-) | INVELSTK | gi\|15022507 +6 | 1.0e5 | 12/14 | 3712 |
| (-) | RINVELSTK | gi\|15022507 +6 | 1.7e4 | 14/16 | 3562 |
| (-) | ASYDDPYKK | gi\|15022507 +4 | 5.9e3 | 13/16 | 2875 |
| (-) | AIEALHGHELRPGR | gi\|15022507 +6 | 2.4e4 | 16/26 | 3582 |
| (-) | LAELSDYRR | gi\|15022507 +3 | 4.5e4 | 13/16 | 3567 |
| (-) | DYAFVHM*EK | gi\|15022507 +6 | 4.4e4 | 12/16 | 3573-3845 |
| (-) | LSESQLSFR | gi\|15022507 +3 | 5.7e3 | 12/16 | 4098 |
| (-) | PRPLNTWK | gi\|15022507 +7 | 1.6e4 | 10/14 | 3474 |
| (-) | ALVVEMSR | gi\|15022507 +7 | 1.4e4 | 12/14 | 3849 |
| (-) | LSESQLSFR | gi\|15022507 +3 | 1.6e4 | 12/16 | 4203-12642 |
| (-) | TQSSASLAASYAAQQHPQAAASYR | gi\|15022507 +5 | 8.0e3 | 24/92 | 3959 |
| (-) | LAELSDYR | gi\|15022507 +3 | 3.1e3 | 9/14 | 3760 |
| (-) | AAIAQLNGK | gi\|15022507 +5 | 4.5e3 | 10/16 | 3247 |
| (-) | INVELSTK | gi\|15022507 +6 | 3.6e3 | 9/14 | 3668 |
| (-) | YSGSYNDYLR | gi\|15022507 +3 | 2.2e3 | 8/18 | 3954 |
| (-) | QPTPPFFGR | gi\|15022507 +5 | 4.8e4 | 9/16 | 4482 |

| | | |
|---|---|---|
| **Ion Current (Sum):** | 1.5e7 | A |
| **Ion Current (Avg):** | 6.7e5 | A B |
| **# MS/MS Spectra:** | 519 | A G |

**A** gi|14141152 **MS/MS Spectra:** 89 **Sum TIC:** 1.0e7 **Avg TIC:** 1.1e5

heterogeneous nuclear ribonucleoprotein M isoform a; heterogenous nuclear ribonucleoprotein M4; N-acetylglucosamine receptor 1; M4 protein; heterogenous nuclear ribonucleoprotein M [Homo sapiens] gi|13111793|gb|AAH00138.2| Heterogeneous nuclear ribonucleoprotein M, isof

| | Sequence | Reference | TIC | Ions | Scan |
|---|---|---|---|---|---|
| (-) | LKEVFSMAGVVVR | gi\|14141152 +11 | 5.8e4 | 22/24 | 5296-5328 |
| (-) | VGEVTYVELLMDAEGK | gi\|14141152 +9 | 7.1e4 | 22/30 | 5824-6273 |
| (-) | EKVGEVTYVELLMDAEGK | gi\|14141152 +9 | 4.4e3 | 23/34 | 6041 |
| (-) | LKEVFSM*AGVVVR | gi\|14141152 +11 | 7.4e4 | 20/24 | 4480-5984 |
| (-) | GIGMGNIGPAGMGMEGIGFGINK | gi\|14141152 +8 | 1.1e4 | 23/44 | 5999 |
| (-) | MEEESGAPGVPSGNGAPGPK | gi\|14141152 +4 | 8.6e4 | 26/38 | 3554-3560 |
| (-) | M*EEESGAPGVPSGNGAPGPK | gi\|14141152 +4 | 1.5e4 | 25/38 | 3549 |
| (-) | M*GLAM*GGGGGASFDR | gi\|14141152 +8 | 3.0e4 | 23/28 | 3611 |
| (-) | EKVGEVTYVELLM*DAEGK | gi\|14141152 +9 | 5.8e3 | 21/34 | 5649 |
| (-) | M*GLAMGGGGGASFDR | gi\|14141152 +8 | 2.5e5 | 22/28 | 3629-9635 |
| (-) | LGSTVFVANLDYK | gi\|14141152 +11 | 3.4e5 | 19/24 | 5158-5690 |
| (-) | VGEVTYVELLM*DAEGK | gi\|14141152 +9 | 8.6e4 | 21/30 | 5301-5990 |
| (-) | AFITNIPFDVK | gi\|14141152 +12 | 1.3e6 | 17/20 | 5571-6705 |
| (-) | INEILSNALK | gi\|14141152 +13 | 5.9e5 | 16/18 | 4488-4883 |
| (-) | GNFGGSFAGSFGGAGGHAPGVAR | gi\|14141152 +13 | 5.8e4 | 19/44 | 4704-4734 |
| (-) | EVFSMAGVVVR | gi\|14141152 +11 | 2.4e5 | 16/20 | 5188-5279 |
| (-) | GIGMGNIGPAGM*GMEGIGFGINK | gi\|14141152 +8 | 3.0e4 | 34/88 | 5601-5860 |
| (-) | M*GAGLGHGMDR | gi\|14141152 +13 | 2.7e4 | 18/20 | 3193 |
| (-) | GIGMGNIGPAGM*GM*EGIGFGINK | gi\|14141152 +8 | 1.2e5 | 24/44 | 5348-5629 |
| (-) | INEILSNALKR | gi\|14141152 +13 | 3.0e4 | 18/20 | 4285-4543 |
| (-) | GEGERPAQNEK | gi\|14141152 +6 | 8.7e3 | 16/20 | 1748 |
| (-) | MGLAMGGGGGASFDR | gi\|14141152 +8 | 2.3e5 | 21/28 | 3994-4419 |
| (-) | GIGMGNIGPAGM*GMEGIGFGINK | gi\|14141152 +8 | 5.6e4 | 22/44 | 5608-5738 |
| (-) | ADILEDKDGK | gi\|14141152 +11 | 2.1e5 | 16/18 | 3061-3318 |
| (-) | QGGGGGGGSVPGIER | gi\|14141152 +8 | 3.0e4 | 20/28 | 3493 |
| (-) | LGGAGMER | gi\|14141152 +8 | 1.1e5 | 13/14 | 2663-3361 |
| (-) | VKEDPDGEHAR | gi\|14141152 +8 | 5.7e4 | 17/20 | 991-2125 |
| (-) | MGAGLGHGMDR | gi\|14141152 +13 | 9.7e3 | 16/20 | 3475 |
| (-) | MGPLGLDHMASSIER | gi\|14141152 +14 | 1.0e5 | 22/28 | 4604-5146 |
| (-) | GGNRFEPYANPTK | gi\|14141152 +8 | 4.2e4 | 16/24 | 3659 |
| (-) | MGQTMER | gi\|14141152 +14 | 1.1e4 | 12/12 | 2433 |
| (-) | KACQIFVR | gi\|14141152 +14 | 1.6e5 | 12/14 | 3663 |
| (-) | GNFGGSFAGSFGGAGGHAPGVAR | gi\|14141152 +13 | 8.3e5 | 30/88 | 4733-12339 |
| (-) | GCAVVEFK | gi\|14141152 +10 | 5.8e4 | 12/14 | 3784 |
| (-) | M*GPAM*GPALGAGIER | gi\|14141152 +13 | 8.7e4 | 17/28 | 3925-4321 |
| (-) | KAAEVLNK | gi\|14141152 +10 | 1.2e5 | 13/14 | 341-9863 |
| (-) | LKEVFSMAGVVVR | gi\|14141152 +11 | 1.6e4 | 26/48 | 5290-5864 |
| (-) | MGAGMGFGLER | gi\|14141152 +14 | 1.7e4 | 15/20 | 4603 |
| (-) | M*GANSLER | gi\|14141152 +27 | 1.3e4 | 12/14 | 2290 |
| (-) | VGQTIER | gi\|14141152 +14 | 1.6e5 | 11/12 | 1958-2549 |
| (-) | VKEDPDGEHAR | gi\|14141152 +8 | 1.2e5 | 25/40 | 22-9548 |
| (-) | M*GAGLGHGMDR | gi\|14141152 +13 | 7.0e4 | 16/20 | 2385-3203 |
| (-) | MVPAGMGAGLER | gi\|14141152 +9 | 2.0e5 | 17/22 | 4018-4209 |
| (-) | EVFSM*AGVVVR | gi\|14141152 +11 | 9.6e4 | 14/20 | 4248-9636 |
| (-) | M*GPAMGPALGAGIER | gi\|14141152 +13 | 2.7e4 | 26/56 | 4058-4449 |
| (-) | MGPAM*GPALGAGIER | gi\|14141152 +13 | 6.3e5 | 18/28 | 4031-4573 |
| (-) | RGGNRFEPYANPTK | gi\|14141152 +8 | 2.0e4 | 16/26 | 3593 |
| (-) | M*GANNLER | gi\|14141152 +14 | 8.4e4 | 12/14 | 1920-3160 |
| (-) | VGEVTYVELLM*DAEGK | gi\|14141152 +9 | 2.4e4 | 21/60 | 5745-6156 |
| (-) | ADILEDKDGK | gi\|14141152 +11 | 1.0e3 | 13/18 | 3289 |
| (-) | DKFNECGHVLYADIK | gi\|14141152 +13 | 7.5e5 | 23/56 | 4381-4405 |
| (-) | MGANNLER | gi\|14141152 +14 | 4.2e3 | 12/14 | 3226 |
| (-) | M*GAGM*GFGLER | gi\|14141152 +14 | 3.6e4 | 13/20 | 3908-4210 |
| (-) | EKVGEVTYVELLM*DAEGK | gi\|14141152 +9 | 8.9e3 | 26/68 | 5639 |
| (-) | M*GLVM*DR | gi\|14141152 +13 | 2.8e4 | 10/12 | 924-8598 |
| (-) | GCAVVEFK | gi\|14141152 +10 | 1.4e5 | 11/14 | 3788-3801 |
| (-) | FGSGMNMGR | gi\|14141152 +14 | 3.5e4 | 11/16 | 3623 |
| (-) | M*GPLGLDHMASSIER | gi\|14141152 +14 | 1.2e5 | 19/28 | 4089-4958 |
| (-) | VGSEIER | gi\|14141152 +13 | 2.3e5 | 11/12 | 2404-3230 |
| (-) | M*GPLGLDHM*ASSIER | gi\|14141152 +14 | 9.7e4 | 25/56 | 3985-4816 |
| (-) | M*GAGM*GFGLER | gi\|14141152 +14 | 1.6e4 | 16/20 | 4318 |
| (-) | MGPAMGPALGAGIER | gi\|14141152 +13 | 1.2e5 | 15/28 | 4421-4678 |
| (-) | M*GANSLER | gi\|14141152 +27 | 6.1e4 | 10/14 | 1985-9933 |
| (-) | M*GPVMDR | gi\|14141152 +14 | 1.6e4 | 10/12 | 2500-3329 |
| (-) | M*GPLGLDHMASSIER | gi\|14141152 +14 | 5.8e5 | 21/56 | 4063-4991 |
| (-) | INEILSNALKR | gi\|14141152 +13 | 2.4e4 | 18/40 | 4263-4490 |
| (-) | HSLSGRPLK | gi\|14141152 +9 | 5.9e4 | 11/16 | 2598-8015 |
| (-) | LGSTVFVANLDYK | gi\|14141152 +11 | 9.6e2 | 14/24 | 12272 |
| (-) | FGSGMNM*GR | gi\|14141152 +14 | 9.6e4 | 10/16 | 3025-3550 |

| | | | | | |
|---|---|---|---|---|---|
| (-) | MGGM*EGPFGGGM*ENMGR | gi\|14141152 +8 | 2.0e5 | 15/32 | 3901-4550 |
| (-) | AAEVLNK | gi\|14141152 +10 | 1.2e5 | 8/12 | 1938-2643 |
| (-) | FESPEVAER | gi\|14141152 +15 | 1.2e4 | 9/16 | 3595 |
| (-) | LGSTVFVANLDYK | gi\|14141152 +11 | 1.2e3 | 12/24 | 5180 |
| (-) | NLPFDFTWK | gi\|14141152 +14 | 8.6e3 | 10/16 | 6015 |
| (-) | AFITNIPFDVK | gi\|14141152 +12 | 4.4e3 | 12/20 | 5658 |
| (-) | M*GLAMGGGGGASFDR | gi\|14141152 +8 | 4.3e3 | 15/28 | 4909 |
| (-) | LGSTVFVANLDYK | gi\|14141152 +11 | 4.2e3 | 11/24 | 5948 |
| (-) | M*VPAGM*GAGLER | gi\|14141152 +9 | 4.2e4 | 14/22 | 3413-3871 |
| (-) | LKEVFSM*AGVVVR | gi\|14141152 +11 | 2.7e3 | 12/24 | 4981 |
| (-) | MGGMEGPFGGGM*ENMGR | gi\|14141152 +8 | 1.4e5 | 17/32 | 4208-4799 |
| (-) | VGQTIER | gi\|14141152 +14 | 6.6e4 | 8/12 | 2379-2508 |
| (-) | MGANSLER | gi\|14141152 +27 | 3.0e3 | 9/14 | 3264 |
| (-) | MGAGMGFGLER | gi\|14141152 +14 | 2.2e4 | 12/20 | 890-9556 |
| (-) | KAAEVLNK | gi\|14141152 +10 | 2.2e3 | 8/14 | 2408 |
| (-) | QGGGGGGGSVPGIER | gi\|14141152 +8 | 3.3e3 | 13/28 | 3514 |
| (-) | KAAEVLNK | gi\|14141152 +10 | 8.3e2 | 8/14 | 2228 |
| (-) | VGSEIER | gi\|14141152 +13 | 8.4e4 | 7/12 | 2419-2809 |
| (-) | AFITNIPFDVK | gi\|14141152 +12 | 9.0e2 | 11/20 | 7041 |
| (-) | INEILSNALK | gi\|14141152 +13 | 4.0e3 | 10/18 | 4643 |

**Supplementary Table S1. Mass spectrometry identification of hnRNP M as CoAA-interacting proteins.** Recombinant CoAA YxxQ domain (307-545) was incubated with HeLa cell nuclear extracts. The two bound protein bands shown in Supplementary Figure S2b were subjected to mass spectrometry analysis. MS/MS spectra of peptide fragments, 102 peptides for the upper band and 89 peptides for the lower band, were analyzed at Harvard Microchemistry and Proteomic Analysis Facility and were identified as hnRNP M. Contaminating recombinant CoAA (coactivator activator) in the gel can also be detected.

**Supplementary Table S2. Primer pairs of the CoAA gene used in ChIP analysis.** The 14 primer pairs for the ChIP analysis (Fig. 3d) are depicted on the human CoAA gene (drawn to scale).

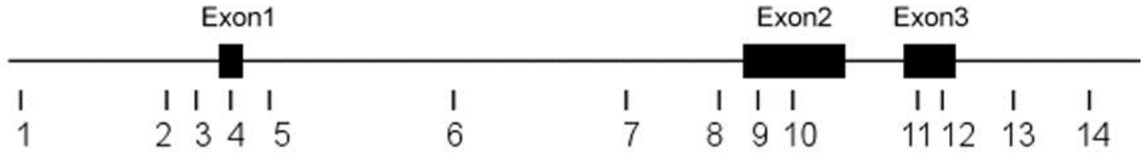


| Primers | Forward primers | Reverse primers |
|---|---|---|
| 1 | AAA CCA AAC CAA ACC AA | GCA ACC CTC TTC GCT CCT |
| 2 | TTA CTC CAA GTC CTC ATC ACC A | TGA CAG AAA TCC CAA CCT TC |
| 3 | CCA GAG ACC GCC TTA CAG AC | AAA CGC TGA GGG AAG ATG TG |
| 4 | GCG GCG ACA AAA TGA AGA TA | TGC ACG AAG GCG AAC TGT |
| 5 | GTC TTG TCT GGC ATG GGT CT | CCC AAA GAA AAA CCC CAA GT |
| 6 | TGG AAC TCA GGC TTC TTG CT | ACT TCA CCC GCA TCG TCA C |
| 7 | AGG CCA AAG GAA ACA CGT AG | AAG GAT GAA ACT CCC TCT C |
| 8 | AAT TGG ACA TGG GCT GAG AG | CCA GCA GAA CAA AAT GCA GA |
| 9 | AAA GGG GAG AAT GGG AAG AA | GCA GCA GAC AGA GGG TTA GC |
| 10 | GCA AAG AAG TGA AGG GCA AG | AAA GCC TGC TGG TAG TCG AA |
| 11 | ATT CCG ATT ACG CAC GCT AT | AGG GCA TGA AAA ACC ACA AC |
| 12 | ACT TTG TTC CTT CGC CTC AG | CCC ATC CAG ACG ACC TTA AA |
| 13 | AAG CAT GGG AGG AGG ATT TT | CAG GAA CCA AGG GAT TGA GA |
| 14 | AGG CTG AAG CAA GAG GAT TG | CCT GCC TCC CAG ACA ATA AA |
